# Single-cell profiling of the human primary motor cortex in ALS and FTLD

**DOI:** 10.1101/2021.07.07.451374

**Authors:** S. Sebastian Pineda, Hyeseung Lee, Brent E. Fitzwalter, Shahin Mohammadi, Luc J. Pregent, Mahammad E. Gardashli, Julio Mantero, Erica Engelberg-Cook, Mariely DeJesus-Hernandez, Marka van Blitterswijk, Cyril Pottier, Rosa Rademakers, Bjorn Oskarsson, Jaimin S. Shah, Ronald C. Petersen, Neill R. Graff-Radford, Bradley F. Boeve, David S. Knopman, Keith A. Josephs, Michael DeTure, Melissa E. Murray, Dennis W. Dickson, Myriam Heiman, Veronique V. Belzil, Manolis Kellis

## Abstract

Amyotrophic lateral sclerosis (ALS) and frontotemporal lobar degeneration (FTLD) are two devastating and fatal neurodegenerative conditions. While distinct, they share many clinical, genetic, and pathological characteristics^1^, and both show selective vulnerability of layer 5b extratelencephalic-projecting cortical populations, including Betz cells in ALS^2,3^ and von Economo neurons (VENs) in FTLD^4,5^. Here, we report the first high resolution single-cell atlas of the human primary motor cortex (MCX) and its transcriptional alterations in ALS and FTLD across ~380,000 nuclei from 64 individuals, including 17 control samples and 47 sporadic and *C9orf72*-associated ALS and FTLD patient samples. We identify 46 transcriptionally distinct cellular subtypes including two Betz-cell subtypes, and we observe a previously unappreciated molecular similarity between Betz cells and VENs of the prefrontal cortex (PFC) and frontal insula. Many of the dysregulated genes and pathways are shared across excitatory neurons, including stress response, ribosome function, oxidative phosphorylation, synaptic vesicle cycle, endoplasmic reticulum protein processing, and autophagy. Betz cells and *SCN4B*+ long-range projecting L3/L5 cells are the most transcriptionally affected in both ALS and FTLD. Lastly, we find that the VEN/Betz cell-enriched transcription factor, POU3F1, has altered subcellular localization, co-localizes with TDP-43 aggregates, and may represent a cell type-specific vulnerability factor in the Betz cells of ALS and FTLD patient tissues.

## Introduction

The primary motor cortex is essential for voluntary motor control, sending layer 5 intratelencephalic tract and pyramidal tract output projections to modulate the execution of movement^6,7^. It is affected in multiple neurodegenerative conditions, including amyotrophic lateral sclerosis (ALS)^2,3^, the most common motor neuron disorder^8,9^, and the related frontotemporal lobar degeneration (FTLD)^10^, the second-most common early onset dementia^8,9^.

While ALS and FTLD are distinct diagnoses, they show overlapping clinical, genetic, and pathological characteristics, suggesting they are part of the same disease spectrum^1^. Clinically, 50% of patients diagnosed with either ALS or FTLD eventually develop symptoms of the other disease, and 20% of patients ultimately receive both diagnoses^10–14^. Genetically, the most common causative mutation for both ALS and FTLD is a repeat expansion in the C9orf72-SMCR8 complex subunit (*C9orf72*)^15,16,17^, with the same mutation leading to ALS or FTLD, even among mutation carriers of the same family. Pathologically, 95% of ALS and ~50% of FTLD patients show a common signature of cytoplasmic accumulation of transactive response DNA binding protein 43 (TDP-43)^18^, an RNA-binding protein critical for RNA processing^18,19^.

We hypothesized that in addition to their shared clinical, genetic, and pathological characteristics, ALS and FTLD may also share a common mechanistic basis at the molecular and cellular level. Indeed, FTLD shows loss of large fork cells and von Economo neurons (VENs) in layer 5 of anterior cingulate, fronto-insular, and dorsolateral prefrontal cortices^20–23^, and ALS shows loss of pyramidal tract upper motor neurons (UMNs), specifically Betz cells, in layer 5 of the primary motor cortex, which make direct corticospinal connections to spinal motor neurons^2,24,25^. Moreover, VENs share molecular markers with Betz cells, including *CTIP2* and *FEZF2^26^*, and recent studies suggest VENs may be extratelencephalic-projecting neurons^27^ similar to Betz cells. Thus, we sought to characterize the vulnerability of neuronal cell types in both ALS and FTLD, with a particular focus on large extratelencephalic-projecting cortical layer 5 neurons using single-cell profiling.

We generated the first high-resolution single-cell molecular atlas of the human primary motor cortex and examined transcriptional alterations in ALS and FTLD across multiple neuronal, glial, and vascular cell types. We profiled ~380,000 nuclei from 64 individuals, including 17 control samples and 47 sporadic and *C9orf72*-associated ALS and FTLD patient samples. We defined 46 transcriptional subtypes including two subtypes of Betz cells that exhibit remarkably close molecular similarity to frontal insula and PFC VENs. We found that excitatory neurons in ALS and FTLD shared a significant number of differentially expressed genes across cell types and phenotypes. For both disorders and across genotypes, we observed the highest degree of transcriptional dysregulation in the Betz cells, and more surprisingly, the L3/L5 *SCN4B*+ long-range projecting neurons. Many of the dysregulated pathways were also common, including stress response, ribosome function, oxidative phosphorylation, synaptic vesicle cycle, endoplasmic reticulum protein processing, and autophagy. Through co-expression network analysis of Betz cells, we identified co-expressed gene modules and their hub genes, both confirming several with known prior associations with ALS and FTLD, and discovering novel ones. Analysis of one of the novel hub genes, the VEN/Betz-cell enriched gene *POU3F1*, revealed that the subcellular localization of the POU3F1 protein is altered and co-localizes with TDP-43 aggregates in Betz cells of ALS and FTLD brain tissue.

## Results

### Characterization of human primary motor cortex

We first sought to characterize the diversity of cell types and marker genes for both neuronal and non-neuronal cells in the human primary motor cortex. After applying stringent quality control metrics and cell filtering, we report 380,610 single-nucleus profiles across 64 primary motor cortex samples, corresponding to ~6000 post-quality control nuclei per donor (**Fig. 1a**, **Extended data table 1**, see Methods). Compared to previous studies^28^, the number of human cells captured represents a 4-fold increase in cell count, 10-fold increase in the number of individuals, and substantial increase in cell level resolution (**Extended Data Figs. 1 and 2**).

**Figure 1.**
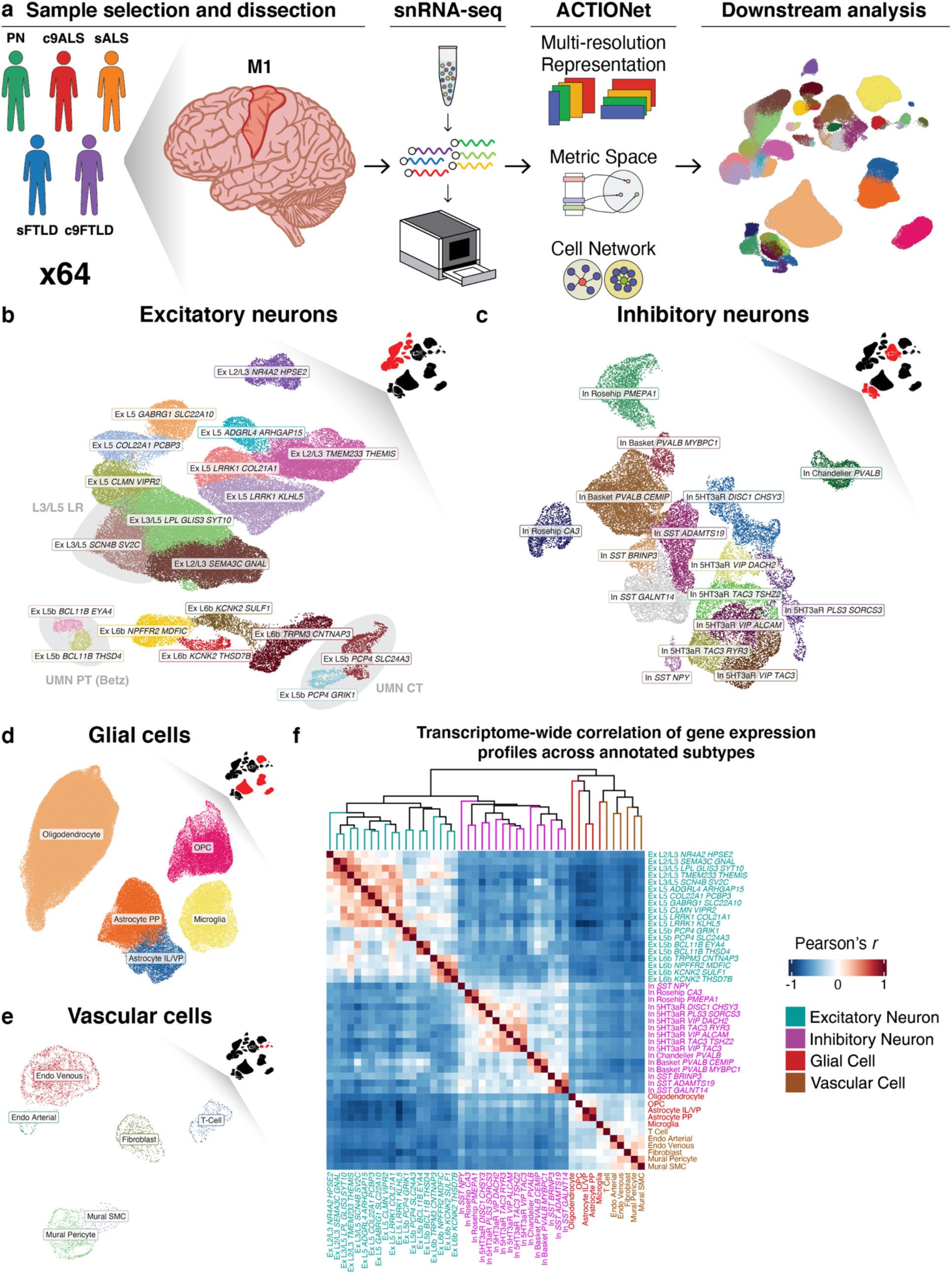
Single-nucleus RNA-seq profiling of primary motor cortex in ALS and FTLD. **a.** Experimental design. **b-e.** Detailed ACTIONet sub-plots of annotated transcriptional subtypes of excitatory neurons (**b**), inhibitory neurons (**c**), glial cells (**d**), and vascular cells (**e**). Insets in upper left corners highlight (in red) clusters of the full ACTIONet plot (**Fig. 1a** right, and **Extended data figure 1a**) shown in each sub-plot. Sub-plot cluster colors correspond to those in the full plot. Clusters of motor and L3/L5 long-range projecting neuron subtypes highlighted in (**b**). **f.** Pairwise Pearson correlation of scaled, mean transcriptome-wide gene counts across annotated subtypes.

We annotated 46 transcriptionally-distinct cell subpopulations (**Fig. 1b–f, Extended Data Figs. 1a and 3**), all of which were well-mixed and reproducible across individuals, sexes, genotypes, and phenotypes (**Extended Data Fig. 1b,d–f**) using our recently-developed single-cell data analysis toolkit, ACTIONet^29^, and well-curated cell type-specific marker genes

We characterized 19 subtypes of excitatory neurons in six major groups (**Fig. 1b**). Translaminar pyramidal neurons spanning layers 2 through 5 formed the largest group, consisting of 6 subtypes based on layer-specific marker gene expression. These showed a gradient of layer-specific marker gene expression, with markers for adjacent layers enriched in adjacent regions of the cluster in a mostly linear fashion, consistent with reports of non-discrete transcriptional identity of translaminar cortical excitatory neurons in mice^30^. No subtype or cluster showed exclusive enrichment for layer 4 markers, consistent with previous reports that layer 4 is absent from agranular primary motor cortex, but adjacent subpopulations with layer 3 and layer 5 markers (Ex L3/L5) and layer 5 subtypes (Ex L5) showed overlapping expression of layer 4 markers. We also detected a distinct subtype of *SCN4B*+ cells expressing markers of both Layer 3 and Layer 5 at the layer 3/5 interface, which showed the highest expression of neurofilament genes *NEFL*, *NEFM*, and *NEFH*, indicative of large axon caliber neurons (**Extended Data Fig. 4c**). This population likely corresponds to *FEZF2*-L3/L5 long-range projecting neurons^31^, and are henceforth referred to as “L3/L5 LR” (**Fig. 1b**).

We identified four distinct subtypes of *FEZF2/CRYM*+ layer 5b neurons partitioned into two groups (**Fig. 1b**; denoted as “UMN PT” and “UMN CT”). The smaller of these two groups corresponded to two transcriptionally distinct subtypes (*BCL11B/EYA4* and *BCL11B/THSD4*) of giant pyramidal tract upper motor neurons (UMN PT), also known as Betz cells, which expressed higher levels of neurofilament gene *NEFH*, and highly-specific enrichment of the Betz and VEN identity marker *POU3F1^32^* (**Extended Data Fig. 4c**). These cells possessed the highest average RNA content among all cell types (**Extended Data Fig. 2a**), indicating a large cell volume characteristic of giant Betz cells. Expression of key markers and expected Betz morphology was confirmed by immunofluorescence (IF) (**Fig. 2a**). These cells are of particular interest as they are known to be selectively vulnerable in ALS^2,33–35^.

**Figure 2.**
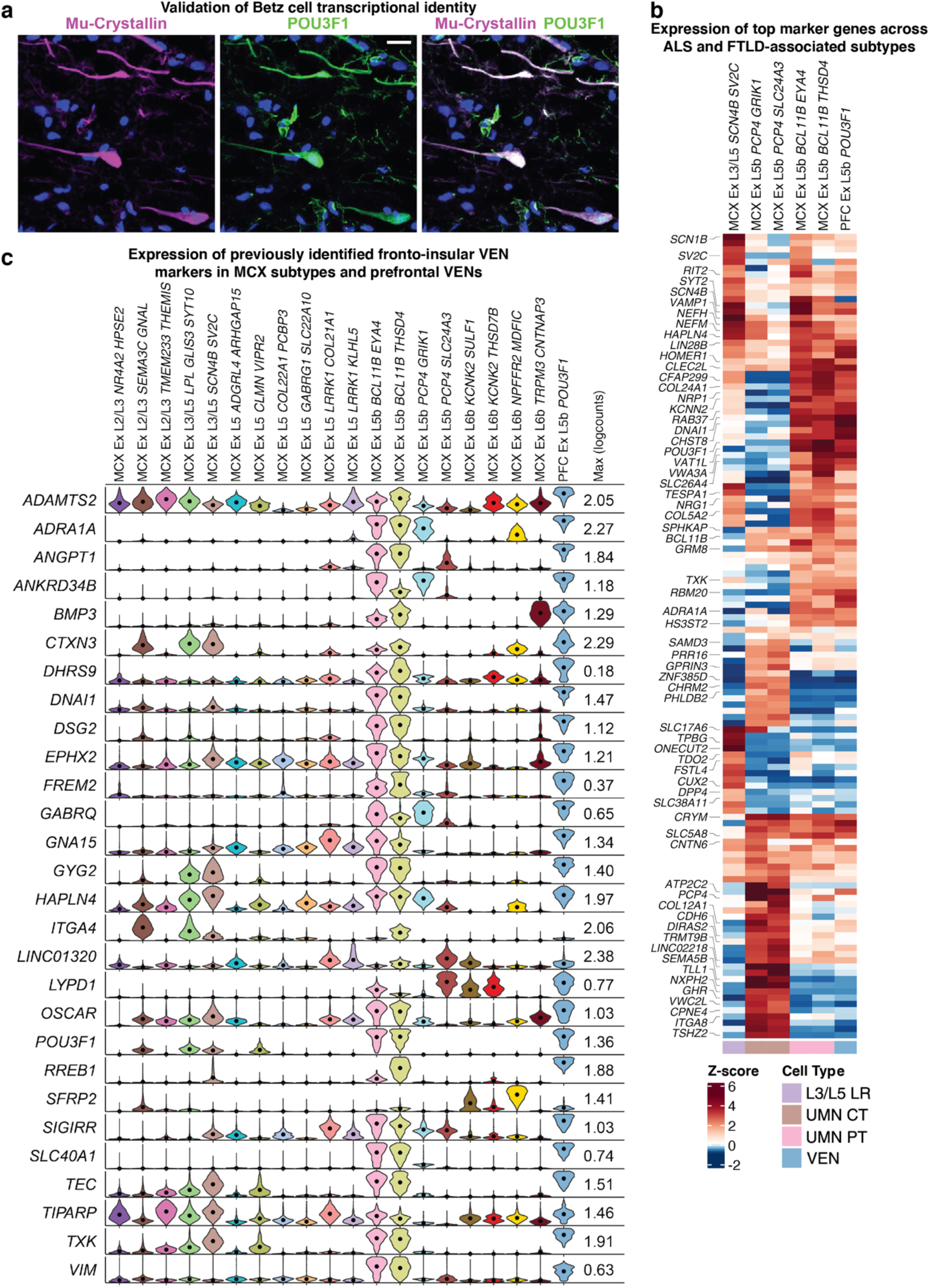
Characterization of ALS and FTLD implicated cell populations. **a.** Representative images showing indirect immunofluorescent labeling of Betz cell markers Mu-Crystallin (*CRYM*) and POU3F1 in pathologically normal tissue. Scale bar = 20μm. **b.** Differential expression of top 50 marker genes of annotated subtypes corresponding to previously described and newly identified differentially vulnerable cell types in ALS and FTLD. Right-most column shows expression of markers in VENs of the dorsolateral PFC. Top 25 markers of each subtype labeled on the left. **c.** Violin plots showing single-cell-level expression of VEN marker genes from Hodge *et al.* (2020) in annotated excitatory subtypes of MCX and VENs of PFC. Both Betz cell clusters in MCX (MCX Ex L5b *BCL11B*) and the VEN cluster in PFC (PFC EX L5b *POU3F1*) exhibit high expression of nearly every identified VEN marker. Dots denote population mean. MCX: motor cortex; PFC: prefrontal cortex.

The larger group of the layer 5b *FEZF2/CRYM*+ cells (*PCP4/GRIK1* and *PCP4/SLC24A3*) was also largely enriched for several canonical UMN markers but was reduced in or lacked expression of the long-range projection markers *NEFH* and *SCN4B* (**Fig. 2b, Extended Data Fig. 4c**). Observation of these cells by IF (not shown) showed that they were also layer 5b pyramidal neurons, but appeared to be much smaller than Betz cells, leading us to conclude that these are likely the corticobulbar tract upper motor neurons (UMN CT) that innervate the cranial nerve nuclei, the reticular formation, and the red nucleus.

We identified a group of four transcriptionally similar excitatory neuron subtypes that showed selective enrichment for markers of layer 6b, and human orthologs of mouse layer 6 subplate-derived deep cortical neurons^36^, as well as a cluster of layer 2/3 neurons with highly-specific expression of *NR4A2* (**Fig. 1b**).

We captured 16 distinct populations of cortical inhibitory neurons (**Fig. 1c**), spanning multiple, highly-resolved subtypes of somatostatin-expressing GABAergic interneurons (*NPY*+ and *NPY*−), parvalbumin-expressing basket and chandelier cells, 5HT3aR-expressing interneurons (*VIP*+ and *VIP*−), and two populations of the recently characterized rosehip interneurons^37^ (*CA3*+ and *PMEPA1*+).

We also recovered all expected classes of cortical glial, vascular, and immune cell types, including oligodendrocytes, oligodendrocyte progenitors, two subtypes (protoplasmic, interlaminar) of astrocytes (**Fig. 1d**), fibroblasts, arterial and venous subtypes of endothelial cells, smooth muscle and pericyte mural cells, and T cells (**Fig. 1e**).

Overall, this dataset represents the largest and most comprehensive transcriptomic characterization of the human primary motor cortex to date. Given the well-curated and reproducible nature of these annotations across individuals, we were able to identify novel markers for all the identified subtypes (**Supplementary Data Table 1**), which will help guide future imaging and other experimental studies to dissect the diverse functional properties and roles of these cell types.

### Motor cortex Betz cells are transcriptional analogues to frontal and temporal cortex VENs

We next asked whether our discovered clusters of Betz cells in the primary motor cortex share intrinsic molecular markers with VENs, motivated by the fact that: (1) Betz cells and VENs display enhanced vulnerability in ALS and FTLD respectively; (2) Betz cells are large extratelencephalic-projecting layer 5 neurons and VENs have been recently hypothesized to also be extratelencephalic-projecting layer 5 neurons^27^.

To compare VENs and Betz cells at similar resolution, we used data from our recent single-cell profiling of the dorsolateral PFC^38^ carried out in the context of schizophrenia, but focusing only on the subset of 24 pathologically normal individuals. Detailed clustering and annotation of 107,358 PFC excitatory neurons (**Extended Data Fig. 4a**) revealed a neurofilament-enriched layer 5b subtype with selective expression of *POU3F1* and co-expression of numerous Betz cell markers (**Extended Data Fig. 4c**), which we determined to be the VEN population.

Pairwise transcriptome-wide analysis revealed that almost all annotated excitatory populations in the MCX could be mapped to a synonymous subtype in the dorsolateral PFC, and that both MCX Betz cell subtypes mapped most specifically to the PFC VEN subtype (*r* = 0.57 and 0.61) (**Extended Data Fig. 4b**).

We then identified and compared the top 50 marker genes of UMN, L3/L5 LR, and VEN subtypes and found that Betz cells and VENs possessed nearly identical expression patterns across these genes, and that a subset of these markers were also highly expressed in L3/L5 LR neurons, but very few were shared with corticobulbar motor neurons (**Fig. 2b**).

When we compared our data with the VEN markers described in a recent, targeted single-cell study of frontal insula VENs^27^, we again found that all reported markers, in addition to the previously described *CTIP2/BCL11B* and *FEZF2^26^*, are also very specifically expressed in Betz cells and, to a lower extent, other layer 5 UMNs and L3/L5 LR cells. This analysis also confirmed the annotation of our PFC VEN cluster (**Fig. 2c**).

This finding reveals a close molecular similarity between the two cortical populations that display the most vulnerability in ALS and FTLD, and provides further evidence for the hypothesis that VENs are in fact extratelencephalic-projecting neurons.

### Cell type-specific differential gene expression analysis of ALS and FTLD

We next performed cell type-specific pseudo-bulk differential expression analysis across all five phenotypes: sporadic ALS (sALS; *n*=17), *C9orf72*-associated ALS (c9ALS; *n*=6), sporadic FTLD (sFTLD; *n*=13), *C9orf72*-associated FTLD (c9FTLD; *n*=11), and pathologically normal (PN; *n*=17) (**Extended Data Table 1**). To ensure that we had sufficient power across all experimental groups for less abundant populations, we aggregated highly similar subtypes within the same group for differential expression analysis (e.g. we treated both transcriptional subtypes of Betz cells as a single population denoted as Ex UMN PT), and excluded In *SST*/*NPY*+ and T cells which were insufficiently abundant across donors and disease groups, and too dissimilar to aggregate with other subtypes (**Extended Data Fig. 5**).

Although large layer 5 projection neurons would be expected to be the most affected in both ALS and FTLD, we expected to see alterations in other cortical layers as well due to spreading of cortical atrophy in late-stage disease. Indeed, we observed a large disparity in the number of dysregulated genes between excitatory neurons and all other cell populations across all phenotypes and found that all excitatory neurons across cortical layers displayed severe transcriptional dysregulation (**Fig. 3a,b, Supplementary Data Tables 2-5**).

**Figure 3.**
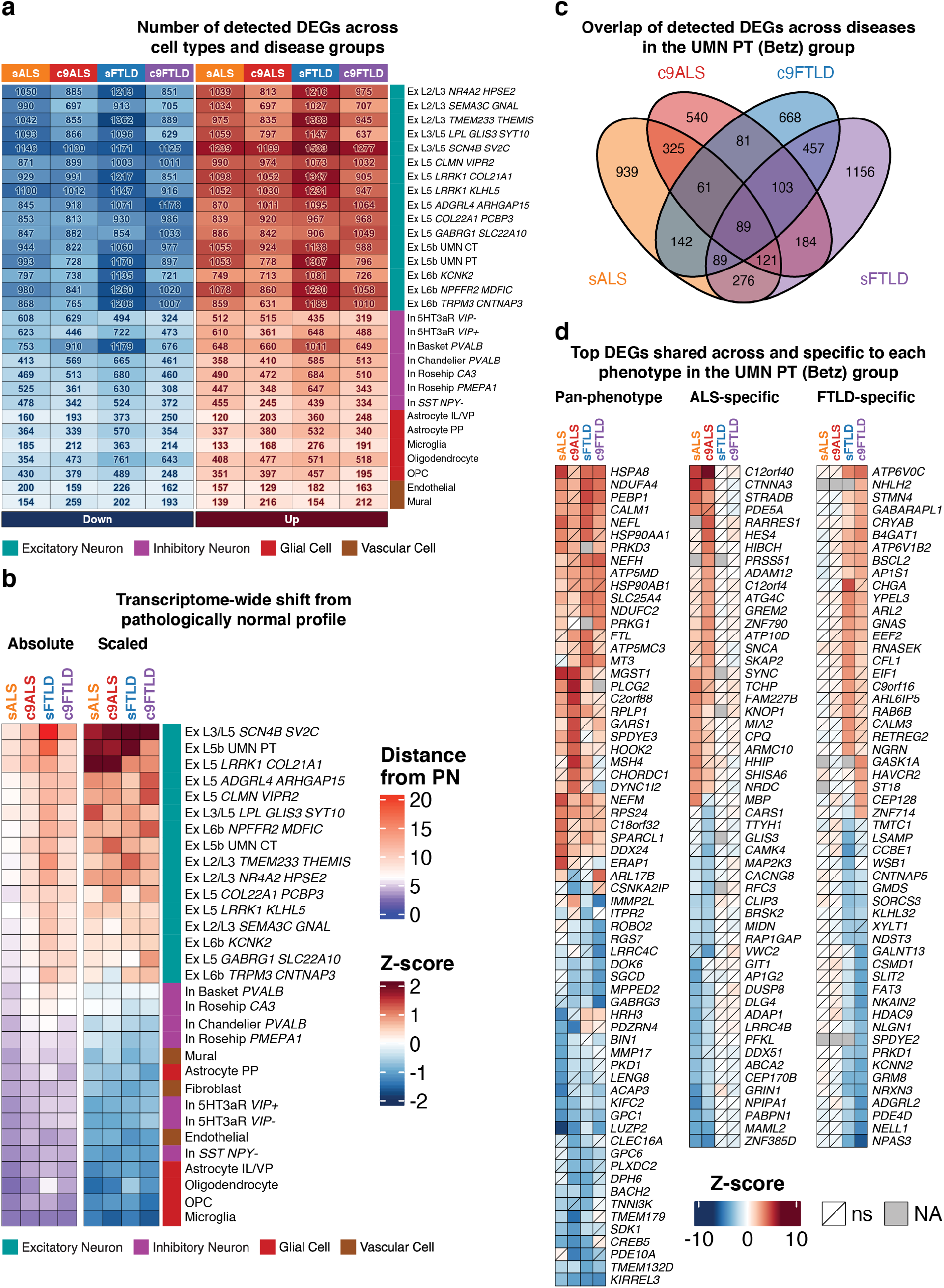
Differential gene expression analysis. **a.** Absolute number of DEGs detected per cell type per disease group (absolute log2-fold change Z-score > 1, FDR-adjusted *p* < 0.005). **b.** Disease score (transcriptomic distance from pathologically normal) of each cell type per disease group. Left: Absolute distance. Right: Column-wise Z-score of distances. **c.** Overlap of detected pyramidal tract upper motor neuron (Betz cell) group DEGs across disease groups. **d.** Top up and down regulated pan-phenotypic and phenotype-specific DEGs (FDR-adjusted *p* < 0.005). ns: Not statistically significant; NA: absent in data set.

Our analysis showed that a significant fraction of upregulated differentially expressed genes (DEGs) was shared across excitatory neurons and appeared across diseases and genotypes (**Extended Data Fig. 6**); most of these genes were differentially expressed only in excitatory neurons.

For example, we observed a pan-phenotypic (that is, present in both ALS and FTLD, either sporadic or *C9orf72*-associated) upregulation of a large number of nuclear-encoded mitochondrial respiratory complex I, III, IV, and V subunit-encoding genes, as well as the mitochondrial membrane transporter ADP/ATP translocase 1 (*SLC25A4*) and the mitochondrial stability regulator mitoregulin (*MTLN*), but not mitochondrial-encoded RNAs, which would be indicative of poor nuclei quality. We also observed a primarily upwards, dysregulation of a substantial fraction of ribosomal subunit-encoding genes belonging to both the cytoplasmic (RPL/S) and mitochondrial (MRPL/S) families, the former of which was dramatically higher in excitatory neurons of c9ALS patients compared to the other cohorts.

Of particular note was the pan-phenotypic upregulation of several other genes previously linked to ALS and FTLD, including several heat shock proteins (*HSP90AA1*, *HSP90AB1*, *HSPA8*), Calmodulin-1 (*CALM1*), and the DEAD-box transcription and splicing regulator *DDX24*, which were among the most highly upregulated genes and present in nearly every excitatory subtype and phenotype and a few inhibitory subtypes (**Extended Data Fig. 6**). As the heat shock response is activated to prevent proteins from denaturing and aggregating under stress conditions, the HSF1 pathway and various heat shock proteins have been studied in the context of ALS and FTLD^39–41^. Indeed Arimoclomol, a co-inducer of the heat shock protein response that may enhance the HSF1 pathway^42–44^, is the focus of multiple clinical trials after delaying disease progression in mice^44^, including a Phase II/III clinical trial for patients with rapidly progressive ALS caused by *SOD1* mutations^45^. Recent results suggest this drug may have therapeutic benefits for FTLD as well^46^. In addition, *CALM1* has been proposed as a potential biomarker of longevity in a *SOD1* mouse model^47^, and *DDX24* was shown to be differentially expressed in ALS blood^48^.

Neurofilament subunits (including *NEFL*, *NEFM*, *NEFH*, *INA*) were another notable class of genes driving the common pan-phenotypic signature. Alterations to *NEFH*, which encodes neurofilament heavy (NHF), are uncommon but found in ALS^49–51^, and overexpression of NFH leads to motor neuron degeneration in transgenic mice^52^. Neuronal cytoskeleton-associated proteins (including *STMN2*, *TUBB2A*, *HOOK2*) were also commonly upregulated genes in excitatory neurons. *STMN2* is highly expressed in the central nervous system^53^, and its expression is increased after neuronal injury^54,55^. Similar to *NEFH,* increased *STMN2* expression is also observed in other neurodegenerative diseases^56–58^ and has been linked to TDP-43 pathobiology. TDP-43 regulates the splicing of *STMN2*, and TDP-43 cytoplasmic re-localization leads to a truncated *STMN2* mRNA, reduced STMN2 protein levels, and reduced neuronal outgrowth^54,59–61^. Even though these changes in cytoskeletal genes were pan-neuronal, their magnitude of upregulation was particularly high in Betz cells for each phenotype (**Fig. 3d**, **Supplementary Data Tables 2-5**), suggesting a possible compensatory response against the degradation of axonal integrity, as Betz cells are most affected. In addition, the L3/L5 LR cell type showed a surprisingly similar, and sometimes more severe, degree of dysregulation of these genes, particularly the neurofilament-encoding genes, indicating that the broad dysregulation of long-range projecting cells is not limited to Betz cells. Lastly, a subtype of L5 cells (Ex L5 *LRRK1 COL21A1*) also showed extensive dysregulation that was particularly prominent in ALS (**Fig. 3b**), but their biological significance is not clear given our current understanding of this population.

Inhibitory neurons showed more heterogeneous disease signatures within and across phenotypes, and significantly less dysregulation overall (**Fig. 3a–b**). Nevertheless, interneuronal subtypes also showed upregulation of heat shock proteins and a few of the top-ranking excitatory DEGs, implying that these genes are part of a pan-neuronal disease or stress response (**Extended Data Fig. 6**). In contrast, all glial and vascular cell types showed little to no overlap in DEGs with either neuronal class or with each other and were overall among the least severely affected cell populations, irrespective of disease.

Across all recovered cell types, downregulated genes showed much more disease-specific patterns, with only a handful of genes appearing across cohorts. However, several genes with links to ALS and FTLD pathobiology were ubiquitously downregulated in neurons within each disease. In both genotypes of ALS, the glutamate receptor subunit-encoding gene *GRIN1*, and the ALS-linked poly(A) binding protein nuclear 1 (*PABPN1*)^62^ were two of the most downregulated genes in most neuronal subtypes. *PABPN1* contains a GCG repeat encoding a polyalanine tract expanded in oculopharyngeal muscular dystrophy (OPMD)^63^, plays a role in the regulation of poly(A) site selection for polyadenylation^64^, interacts with *TDP-43*, *MATR3* and *hnRNPA1^62,65,66^*, three genes mutated in both ALS and FTLD^67–69^, and is a suppressor of TDP-43 toxicity in ALS models^62^. The *C9orf72* gene itself was identified as differentially expressed in a small, non-specific subset of excitatory subtypes, and only in *C9orf72*-associated cohorts, but was only marginally downregulated in these patients.

Since the absolute number of DEGs in a cell population can be biased by various factors (e.g., number of cells, differences in endogenous gene expression across cell types, cellular RNA content), we computed the difference of the mean, transcriptome-wide distance across all cells between diseased and pathologically normal populations for all cell types and disease groups and used this distance as a disease severity score. This analysis revealed that across cell types, both FTLD cohorts showed a much stronger disease signature than ALS cohorts (**Fig. 3b**). It also showed that in all disease subgroups, L3/L5 LR and Betz cells exhibited the most drastic transcriptome-wide shift from the pathologically normal profile, with the former showing more dysregulation than the Betz population in three of the four groups (**Fig. 3b**). This suggests that while ALS- and FTLD-induced transcriptome perturbations may not be unique to Betz cells, these and the ill-defined L3/L5 LR cells, exhibit enhanced vulnerability in both ALS and FTLD relative to other cell types.

The same observation was not made for corticobulbar motor neurons, which showed a degree of transcriptional misregulation comparable to most other non-Betz excitatory neurons. Taken together, our results suggest that, in the primary motor cortex at a molecular level, *SCN4B/SV2C*+ L3/L5 long-range projecting cells (which are *FEZF2*- and were previously hypothesized to be intratelencephalic projecting^31^) and L5 Betz long-range extratelencephalic-projecting cells are the most affected cell types assessed in this study, for both ALS and FTLD. These cells, and to a lesser extent all other excitatory neurons, display dysregulation of several genes recognized to be genetically or mechanistically linked to ALS and FTLD.

Comparison of Betz cell DEGs across sporadic and *C9orf72*-associated samples showed strong agreement within genotypes of both ALS (R = 0.84) and FTLD (R = 0.77), with FTLD showing more heterogeneity (**Extended Data Fig. 7a**). All detected genotype-specific DEGs were uniquely perturbed in either sporadic or *C9orf72*-associated samples, with none of the highly altered genes showing anti-directional dysregulation in either disease (**Fig 3d**, **Extended Data Fig. 7a**). Cross-phenotype comparisons showed weaker, but still respectable correlations, with most uniquely dysregulated or anti-directional perturbations being specific for ALS or FTLD (**Extended Data Fig. 7b**). The marginal differences in transcriptional perturbations observed between genotypes is consistent with the fact that, in the absence of genotype information, sporadic and *C9orf72*-associated cases of ALS and FTLD are clinically indistinguishable. More broadly, a high-level comparison of DEGs across all cell types revealed that, similar to Betz cells, all other cell types showed modest intra-subtype agreement in transcriptional alterations, even across dissimilar disease groups where few or no other subtype pairs exhibited any degree of similarity (**Extended Data Fig. 8**).

### Betz cell biological pathway alterations in ALS and FTLD

As Betz cells exhibited a large transcriptome-wide shift in both ALS and FTLD and are known to be affected in both diseases^10^, we sought to identify trait-specific putative gene regulatory networks that are specifically altered in this cell population. For this purpose, we used weighted gene co-expression network analysis (WGCNA), which summarizes co-expressed gene clusters into “modules” and is widely utilized to identify highly correlated genes via unsupervised clustering^70–72^. Using pseudo-bulk transcriptional data converted from the single-nuclear transcriptional data described above, our analysis revealed 46 modules specific to ALS and FTLD Betz cells (**Fig. 4a**). Among these, 19 modules significantly correlated with clinical traits (sporadic and *C9orf72*-associated ALS and FTLD; **Fig. 4b**). We found that sporadic FTLD had the most correlated modules, followed by c9FTLD, c9ALS, and sALS. Notably, the “darkorange2” module correlated with both sporadic ALS and FTLD groups. Interestingly, all four members of the stathmin family of genes (*STMN1*, *STMN2*, *STMN3*, and *STMN4*) were identified as top hub genes for this module, having expression values highly correlated with the “darkorange2” module’s eigengene values^73^ (**Supplementary Data Table 6**). Many hub genes of the “darkorange2” module were also known ALS- and FTLD-associated genes, including *DCTN1*, *CHCHD10*, *SOD1*, *SQSTM1*, *VAPB*, *VCP*, *UBQLN2*, *PFN1*, and *PRNP*, a known *C9orf72* age-of-onset modifier^74–76^.

**Figure 4.**
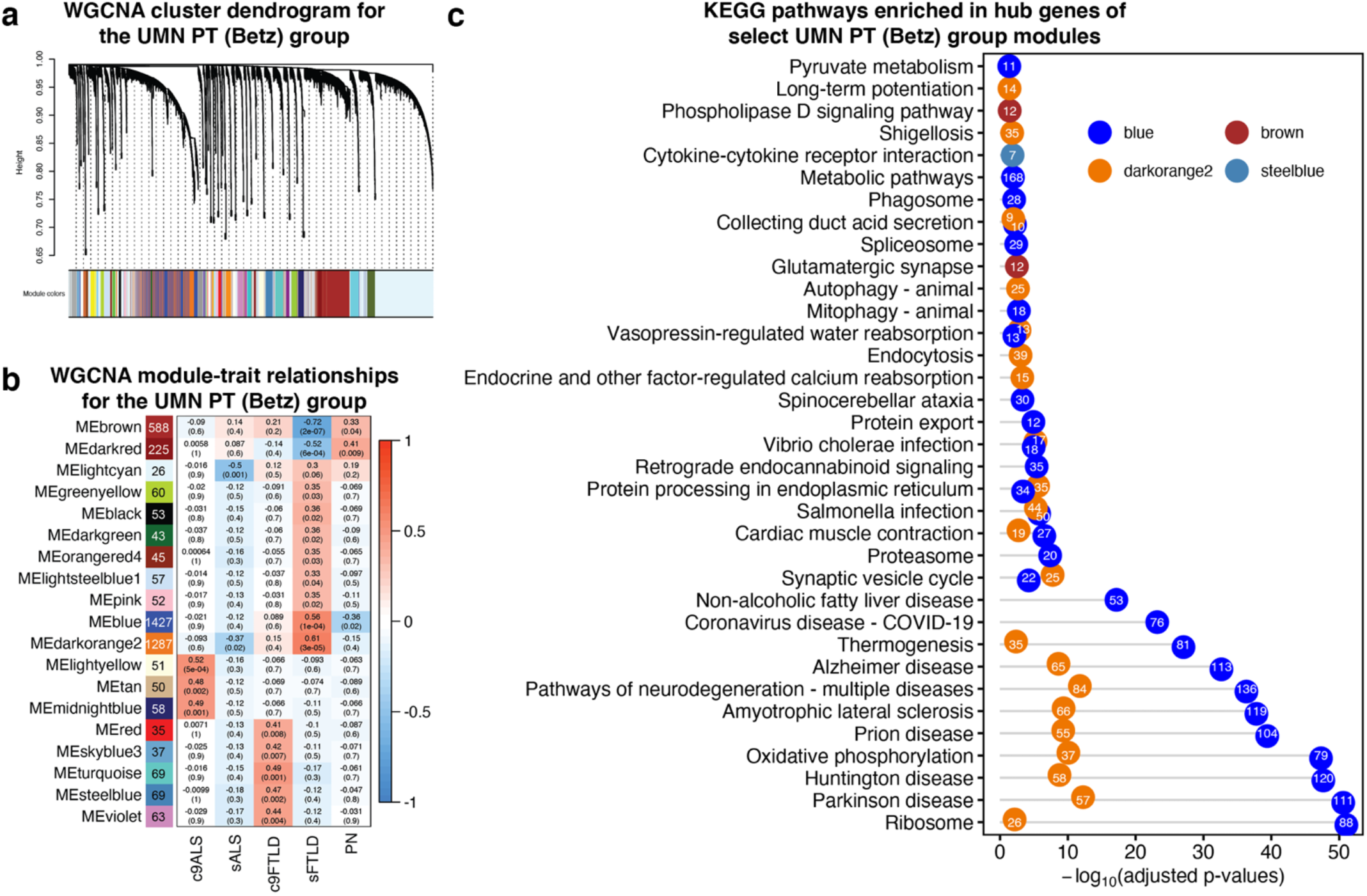
WGCNA analysis of Ex UMN PT (Betz) cluster in ALS and FTLD. **a.** Dendrogram showing result of unsupervised hierarchical clustering of modules identified by WGCNA. **b.** Pearson correlation of module eigengenes with disease groups. Modules significantly correlated with ALS and/or FTLD are shown, with the number of hub genes displayed on the right of each module’s name. **c.** Significant KEGG pathways enriched in hub genes of “blue”, “brown”, “steelblue”, and “darkorange2” modules. Numbers in each circle represent the number of hub genes in each pathway from the corresponding module.

Analysis of hub genes for the 19 modules revealed many shared pathways from both “blue” and “darkorange2” modules (**Fig. 4c**, **Supplementary Data Table 7**). The significant “blue” and “darkorange2” pathways were related to stress response, and included terms such as ribosome, oxidative phosphorylation, synaptic vesicle cycle, protein processing in endoplasmic reticulum, and autophagy. Importantly, impaired stress response is indeed a widely recognized pathological mechanism for both ALS and FTLD, and includes oxidative stress, endoplasmic reticulum stress, disruption of major protein clearance pathways such as ubiquitin-proteasome system and autophagy, altered stress granules dynamics, unfolded protein response, and DNA damage/repair response^77–79^. Also of interest, nuclear pore complex (NPC) and nucleocytoplasmic transport (NCT) defects were common terms for sporadic and *C9orf72*-associated ALS and FTLD, and hub genes included nuclear pore complex *NUP50*, nuclear transport receptor *TNPO3*, and arginine methyltransferase *PRMT1*. NPC and aberrant NCT have received a great deal of interest in recent years after being first observed in *C9orf72*-associated diseases and then sporadic cases^80,81^. The hub genes identified here are also known modifiers of NPC and NCT, and are associated with multiple neurodegenerative diseases including ALS and FTLD^82^, including Nup50 mutations demonstrated to be genetic suppressors of TDP-43 toxicity^83,84^. Disruption in NPC and NCT affects the localization of multiple proteins involved in the stress response such as ribosomal proteins and stress granule-associated RNA binding proteins^84^. A number of hub genes identified encode proteins interacting with G3BP1 (*CAPRIN1*, *CSDE1*, *POLR2B*, *EIF3G*, *DDX3X*, *NUFIP2, EEF2*), which is known to specifically bind to the Ras-GTPase-activating protein, a key regulator of NCT^85,86^. Of interest, dysregulated NPCs were shown to be degraded via upregulation of the ESCRT-III/Vps4 Complex in Drosophila models of *C9orf72* ALS^87^, and components of the four core subunits of the ESCRT-III complex (*CHMP4B*, *CHMP1A*, *CHMP5*) and of other ESCRT complexes (*VPS28* and *VPS25*)^88^ were also “darkorange2” hub genes.

Further inspection of hub genes revealed other interesting regulatory networks that are potentially contributing to ALS and FTLD pathogenesis, such as *SRSF3* and *SRSF8*. SRSF proteins are pre-mRNA splicing factors with multiple functions, including mRNA export from the nucleus^89^. Some SRSF proteins have been found to be sequestered by *C9orf72*-associated RNA foci (SRSF1; SRSF2), and SRSF1, SRSF3, SRSF7 were shown to have increased binding to the GGGGCC (G4C2) *C9orf72* expanded repeat^90^, potentially overriding normal nuclear retention, encouraging nuclear export of repeat-expanded pre-mRNAs, and consequently leading to repeat-associated non-AUG (RAN) translation and dipeptide repeats (DPR) in *C9orf72*-associated diseases^91,92^. Other hub genes included the RNA-G4s helicase *DDX3X*, encoding a protein involved in transcriptional regulation, pre-mRNA splicing, and mRNA export, and part of the DEAD-box protein family characterized by a conserved Asp-Glu-Ala-Asp (DEAD) motif^93^. DDX3X associates with 5’-UTR RNA G-quadruplexes (rG4s)-containing transcripts^94^, which is especially relevant for *C9orf72*-associated ALS and FTLD, as the G_4_C_2_ *C9orf72* expanded repeat leads to a repeat-length-dependent accumulation of rG4-containing transcripts^95^. Notably, DDX3X was shown to directly bind G_4_C_2_ RNAs, consequently suppressing RAN translation, DPR production, and aberrant nucleocytoplasmic transport; its helicase activity is essential for such translation repression^96^. *EIF2AK2*, another hub gene, encodes the protein kinase R (PKR) which plays a key role in mRNA translation, transcriptional control, and regulation of apoptosis^97^, and is regulated by double-stranded RNAs or double-stranded RNA-binding proteins^98^. *TIA1*, a gene mutated in both ALS and FTLD^99^ and a stress granule marker that colocalizes with TDP-43 inclusions in ALS and FTLD^100^, has been demonstrated to be essential for appropriate activation of the PKR-mediated stress response^101^. A direct connection between *EIF2AK2* and TDP-43 was demonstrated when induced TDP-43 toxicity in flies upregulated phosphorylation of *EIF2AK2^102^*. In addition, *MARK2*, a hub gene in the “lightcyan” module, encodes a protein involved in the stability control of microtubules. MARK2 specifically phosphorylates eIF2α in response to proteotoxic stress and is activated via phosphorylation in ALS^103^. Inhibition of eIF2α-phosphorylation has been demonstrated to mitigate TDP-43 toxicity^102^.

We also observed hub genes involved in N6-methyladenosine (m6A) RNA metabolism such as *RBM15B*, a component of a regulatory protein complex that regulates m6A “writer”, and *ALKBH5*, an m6A “eraser” involved in global m6A demethylation. Genes involved in the degradation of m6A-containing mRNAs such as deadenylase *CNOT7* and *RPP25*, a shared component of ribonuclease (RNase) P and RNase mitochondrial RNA-processing (MRP), were also found as hub genes in the “darkorange2” module. These observations indicate potential disruption of the m6A RNA methylome in Betz cells.

Additional WGCNA analysis of the cell type showing the highest transcriptional dysregulation, the L3/L5 LR cells, identified 20 modules significantly correlated with the different clinical traits (**Extended Data Fig. 9**). Pathway analysis of hub genes from these modules revealed many overlapping pathways altered in the Betz cells, suggesting disruptions of similar pathways in both cell types.

Together, our findings highlight various ‘hub’ genes and biological pathways with prior association with ALS and FTLD, reinforcing the notion that transcriptional dysregulation can be used not only as a molecular marker of disease, but also as a resource to identify new candidate driver genes. Considering putative driver genes, of particular interest is the VEN/Betz cell enriched alanine-repeat encoding gene *POU3F1^104^*, which appeared as a top 10% “darkorange2” module hub gene (kME value of 0.87, 97th among 1,287 total hub genes). The observation that *POU3F1* is highly enriched in VENs and Betz cells (**Fig. 2, Extended Data Fig. 4c**), that it contains a GCG repetitive sequence similar to *PABPN1*, and that its dysregulation leads to axonal loss^105^, suggest possible involvement in ALS and FTLD pathobiology.

### POU3F1 Alterations in ALS

As short (8-13 repeats) alanine-encoding GCG expansions in the *PABPN1* gene cause OPMD^63^, which is characterized by TDP-43-positive aggregates^106^, it is possible that the 11-alanine repeat encoding hub gene *POU3F1* represents an intrinsic vulnerability factor of Betz cells and VENs in ALS and FTLD by similarly influencing TDP-43 aggregation. To investigate whether POU3F1 co-localizes with TDP-43 aggregates in ALS and FTLD, we conducted indirect immunofluorescent staining of ALS and FTLD primary motor cortex post-mortem tissue samples (**Fig. 5**). We found that POU3F1 displayed a broad subcellular distribution in Betz cells in pathologically normal tissue. However, in ALS and FTLD patient tissue, we observed a shift in POU3F1 subcellular localization, which now exhibited a punctate morphology in the cytosol of Betz cells. Strikingly, we found that this altered localization of POU3F1 often co-localized with TDP-43 aggregate puncta in Betz cells (**Fig. 5**), suggesting that TDP-43 may co-aggregate with this Betz cell-enriched transcription factor and contribute to cell type-specific dysfunction in ALS and FTLD. Taken together, these results suggest that the Betz/VEN-enriched transcription factor POU3F1 is mislocalized, consequently impairing its normal cellular functions in ALS and FTLD patient Betz cells.

**Figure 5.**
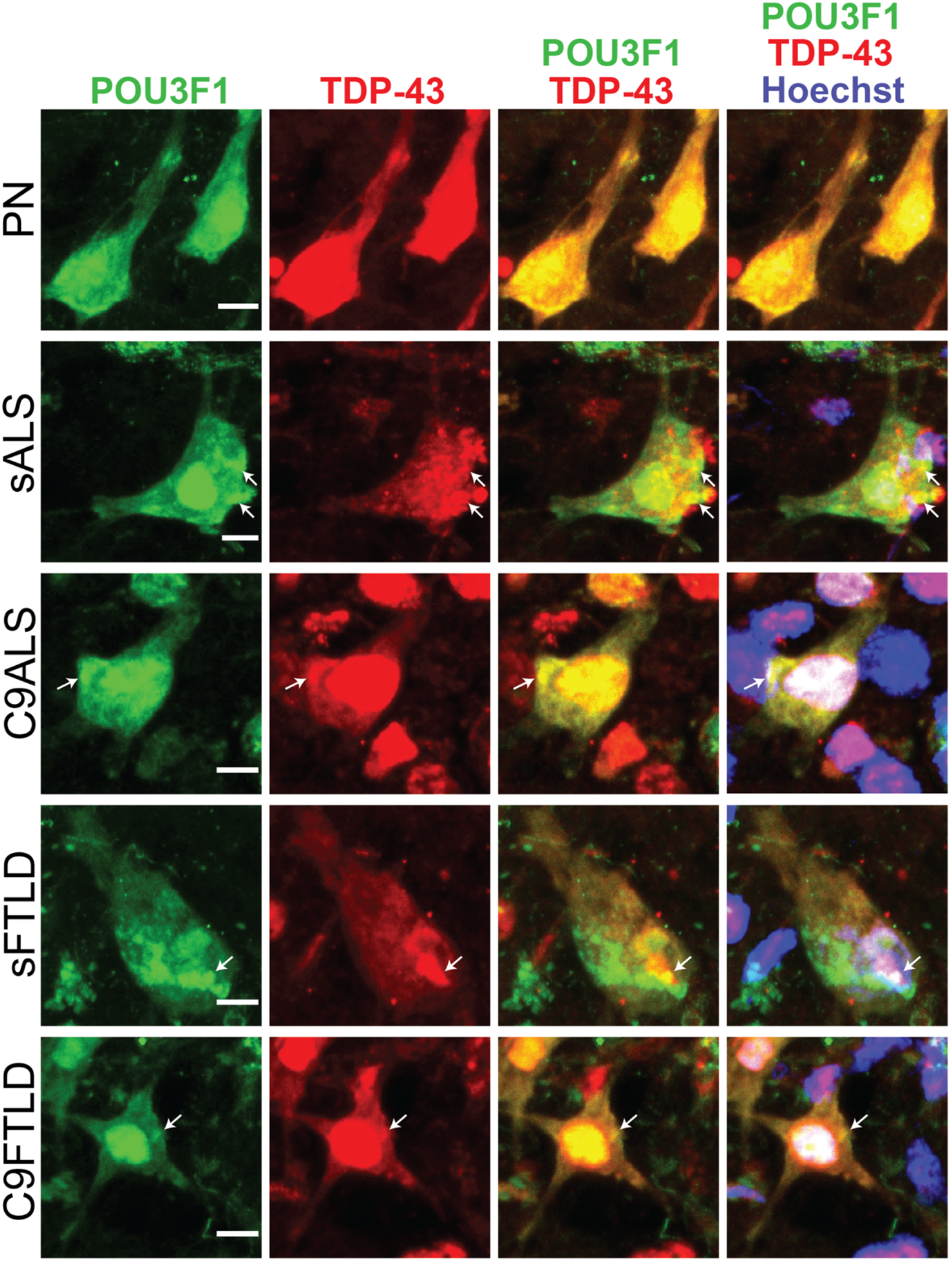
POU3F1 is enriched in CRYM+ cells and displays altered subcellular localization in ALS and FTLD patient tissues. Representative images showing indirect immunofluorescent labeling of POU3F1 and TDP-43 (*TARDBP*) in Betz cells of pathologically normal (PN), sporadic ALS (sALS), *C9orf72*-associated ALS (c9ALS), sporadic FTLD (sFTLD), and *C9orf72*-associated FTLD (c9FTLD) patient tissues. Arrows indicate select regions displaying colocalized, punctate morphology of POU3F1 and TDP-43 fluorescent signals. Scale bar = 10μm.

## Discussion

Our study represents the largest and most comprehensive molecular atlas of the human primary motor cortex, revealing previously unknown characteristics of this brain region, as well as the first documentation of single-cell-level molecular alterations in ALS and FTLD. These will serve as a resource to not only the ALS and FTLD research communities, but also to the human genomics field at large.

We report the existence of at least two distinct classes of Betz cells, as well as a previously unappreciated close molecular similarity between Betz cells of the motor cortex and VENs of the frontal insula and dorsolateral PFC, uncovering a novel link between these vulnerable brain regions and lending further evidence to the notion of an ALS-FTLD pathological spectrum. Further studies will be needed to understand how the two classes of Betz cells differ at a functional level, as well as in the context of ALS and FTLD.

We reveal that, in addition to these Betz cells, a recently identified *SCN4B/SV2C*+ long-range projecting L3/L5 cell type is the most transcriptionally affected in both ALS and FTLD. In the original description^31^, it was postulated that this cell type’s markers represent a shift of expression from preferentially layer 5 in mouse to preferentially layer 3 in human^31^. The authors went on to suggest that this population reflects a unique set of human layer 3 pyramidal neurons that may have human (or primate)-specific long-range intracortical projections^31^. Our studies confirmed that these cells are *FEZF2*-, but that they otherwise express several layer 5 markers (we thus assign them L3/L5 identity), as well as all three neurofilament triplet genes, supporting the idea that they are indeed long-range projection neurons. Further studies will be needed to better understand the cellular and molecular characteristics of this cell population in the normal human brain, as well as in the context of ALS and FTLD, in which they appear to be even more vulnerable than Betz cells. Despite no widespread reports of cell death in this population, some previous pathological studies have suggested that alterations in ALS motor cortex may in fact begin in some pyramidal cells of layer 3^107^. Interestingly, our motor cortex profiles show that these L3/L5 *SCN4B/SV2C*+ cells are second to Betz cells in terms of sharing molecular similarity to VENs (**Fig. 2b,c**), further reinforcing that these intrinsic molecular characteristics, when possessed by cortical neurons, confer vulnerability to cell loss or dysregulation in ALS and FTLD.

Our studies also reveal many ALS and FTLD-associated pathways and likely driver genes, including the VEN/Betz-cell enriched gene *POU3F1*, for which we demonstrated that its encoded protein co-aggregates with TDP-43 in Betz cells of ALS and FTLD brain tissue. *POU3F1* has been ascribed various roles in the developing nervous system, including in neuronal fate commitment^108^, motor neuron identity^32,109^, oligodendrocyte differentiation^110^, and Schwann cell differentiation^111^. Developmental perturbations to *POU3F1* activity result in axonal abnormalities, myelination abnormalities, and premature death, with knockout of *POU3F1* resulting in a fatal breathing defect^105,112,113^. However, *POU3F1* expression patterns change during development, and its role in the adult nervous system is poorly understood, as is its role in ALS and FTLD pathobiology. Future studies will be needed to understand the role of POU3F1 and other markers shared exclusively across Betz, VEN, and L3/L5 LR cells that our results suggest may be key factors underlying the differential vulnerability of these cell types in ALS and FTLD pathogenesis.

---

**Extended Data Figure 1.**
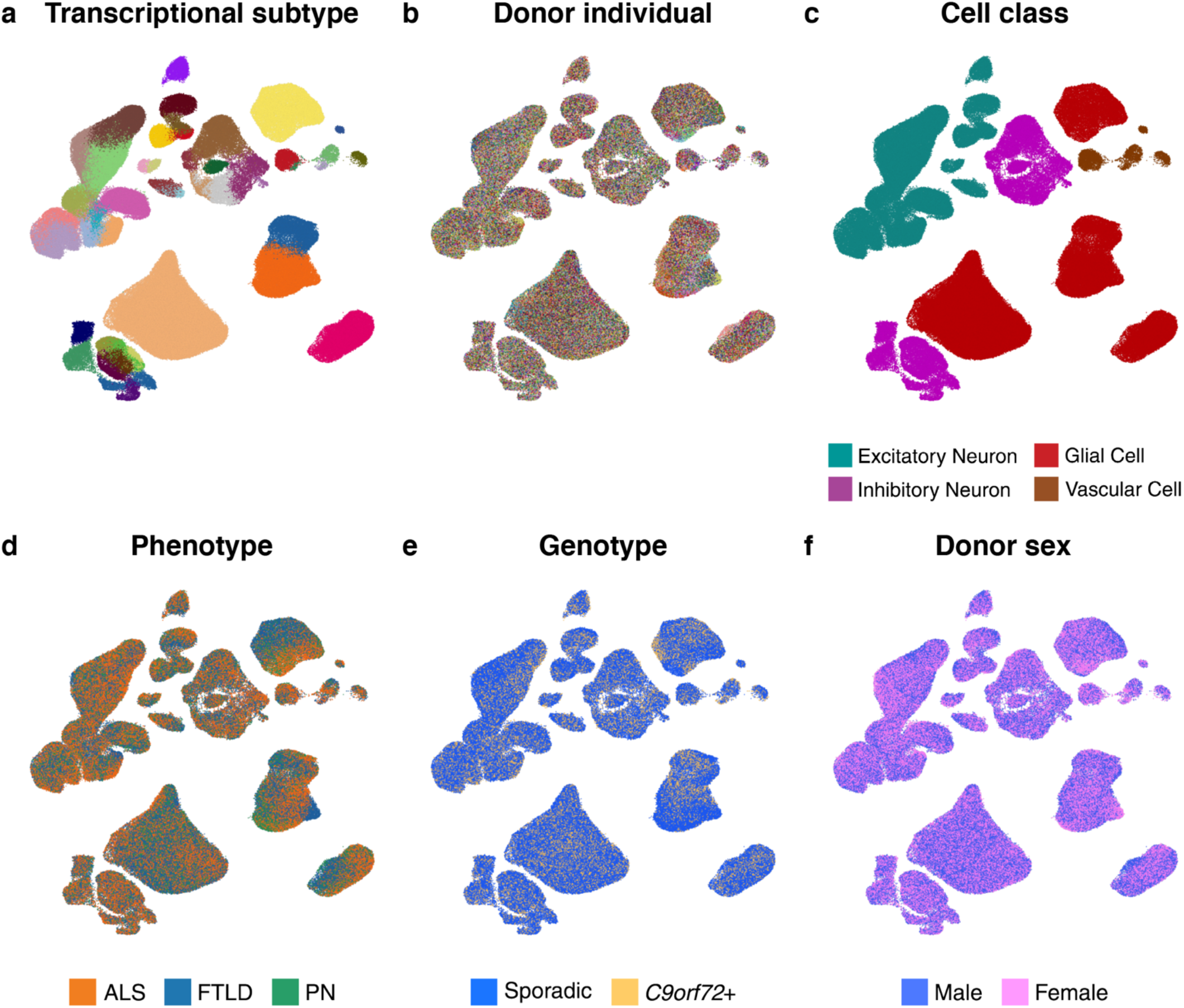
ACTIONet clustering results colored by attribute. **a-f.** Full ACTIONet plots (380,610 cells) colored by transcriptional subtype (**a**), donor individual (**b**), major cell class (**c**), disease phenotype (**d**), disease genotype (**e**), and donor sex (**f**).

**Extended Data Figure 2.**
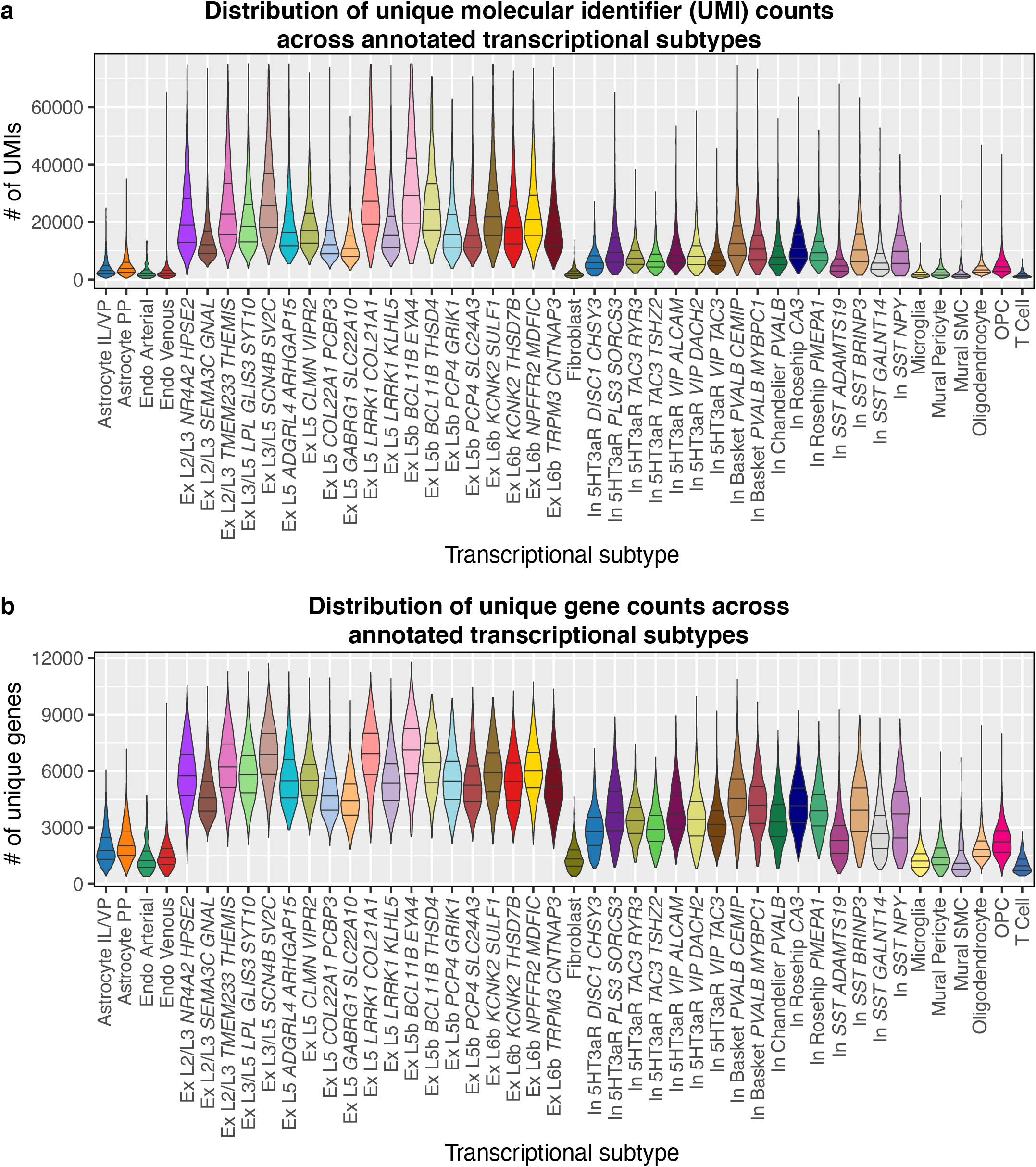
Gene expression complexity across annotated transcriptional subtypes. **a.** Distribution of total unique molecular identifier (UMI) counts per transcriptional subtype. **b.** Distribution of total unique genes detected per transcriptional subtype.

**Extended Data Figure 3.**
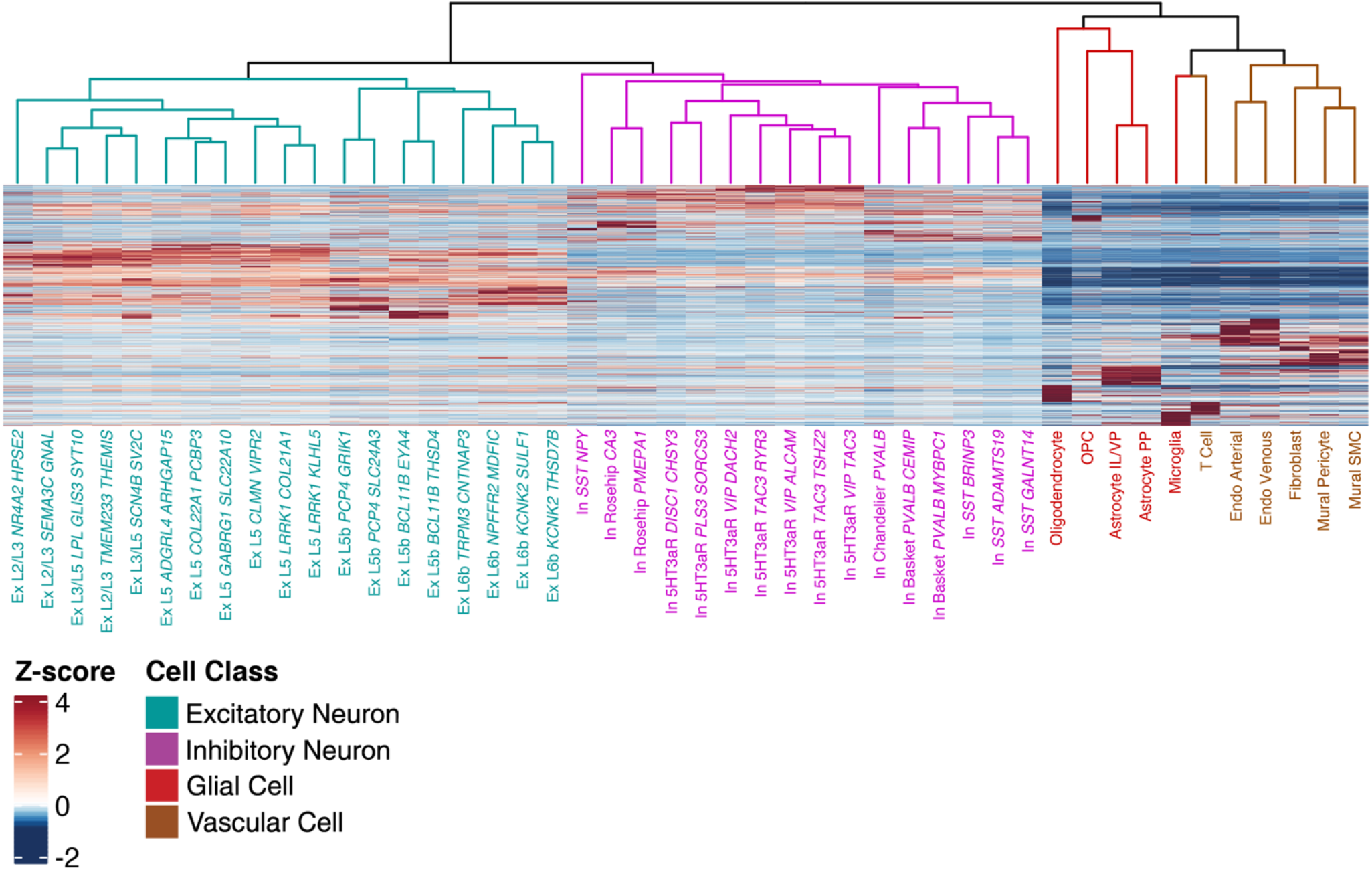
Taxonomy of annotated subtypes. Dendrogram: Hierarchical clustering of transcriptional similarity by mean cluster expression of all genes expressed in > 0.5% of cells (19,953 genes). Identical to dendrogram on left side of **Figure 1f**. Heatmap: Visualization of the expression pattern of the top 200 marker genes per transcriptional subtype (2,683 genes). Color denotes row-wise Z-score of normalized expression.

**Extended Data Figure 4.**
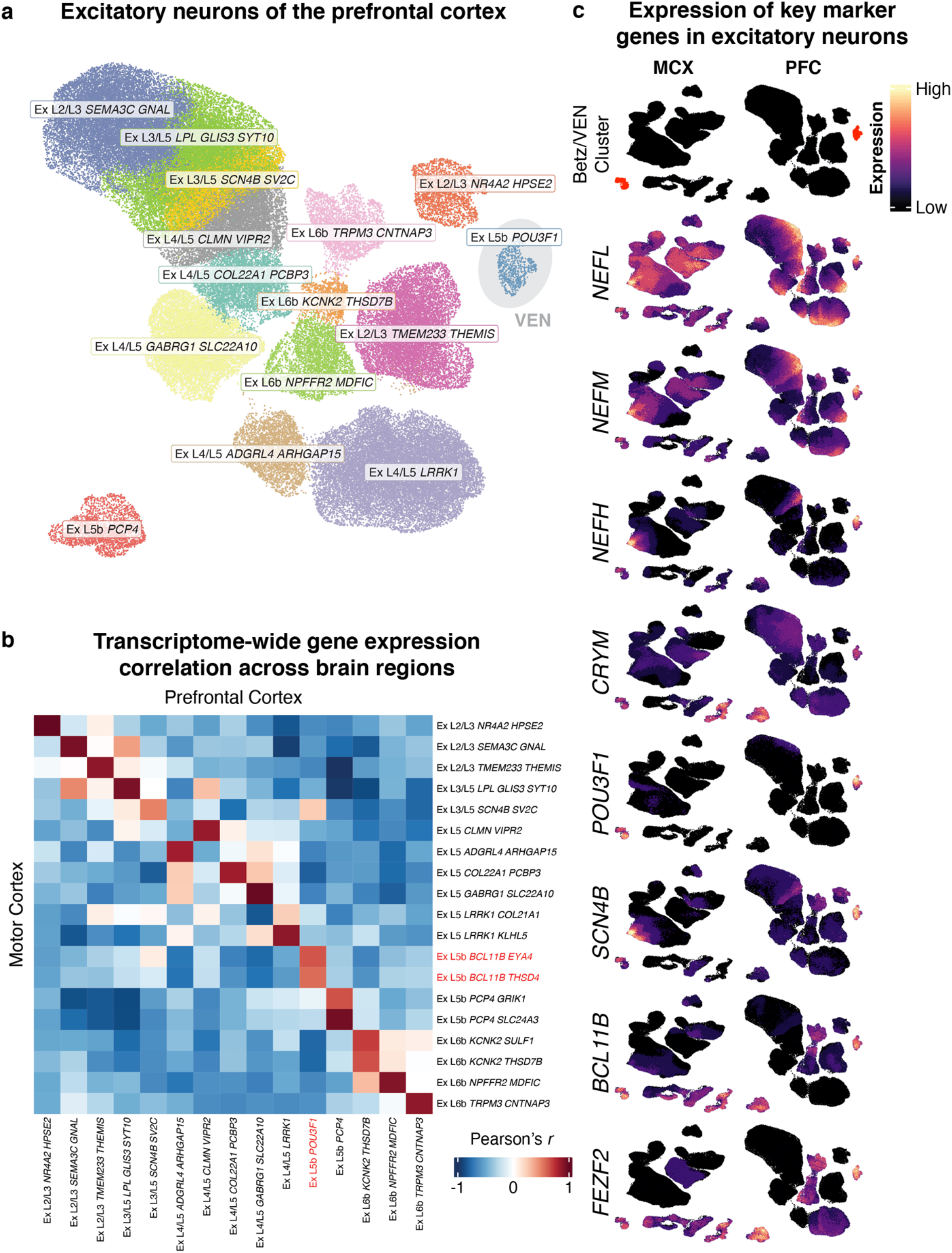
Cross-region comparison of identified excitatory neuron subtypes. **a.** ACTIONet plot of excitatory neurons (107,358 cells) from Ruzicka *et al.* (2020). Cluster identified as von Economo neurons (VENs) highlighted. **b.** Cross-region, pairwise Pearson correlation of normalized, transcriptome-wide gene expression across excitatory subtypes. Names of clusters corresponding to Betz and VEN subtypes highlighted in red. **c.** Relative expression heatmaps of neurofilament subunits and select Betz, VEN, L3/L5 LR, and layer 5b marker genes. Legend (top) denotes clusters (highlighted in red) corresponding to Betz cells (left) and VENs (right).

**Extended Data Figure 5.**
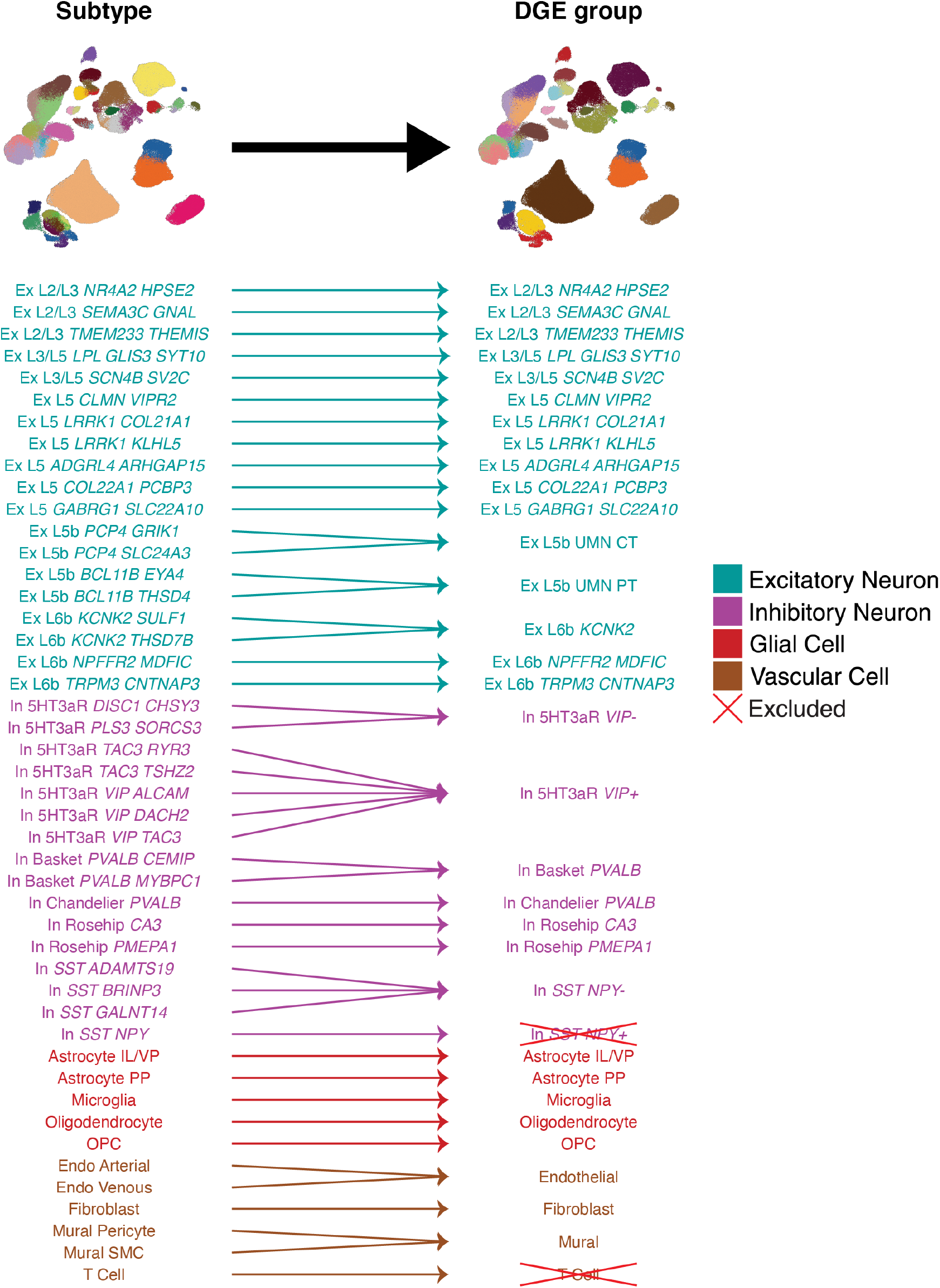
Mapping of annotations of transcriptional subtypes to differential expression analysis group. **a.** Highly similar cell populations of less abundant subtypes were aggregated to ensure sufficient cell counts and donor representation across all disease groups. Arrows show how adjacent subtype clusters were aggregated. Groups marked as “excluded” had insufficient cell counts or donor representation for downstream analyses.

**Extended Data Figure 6.**
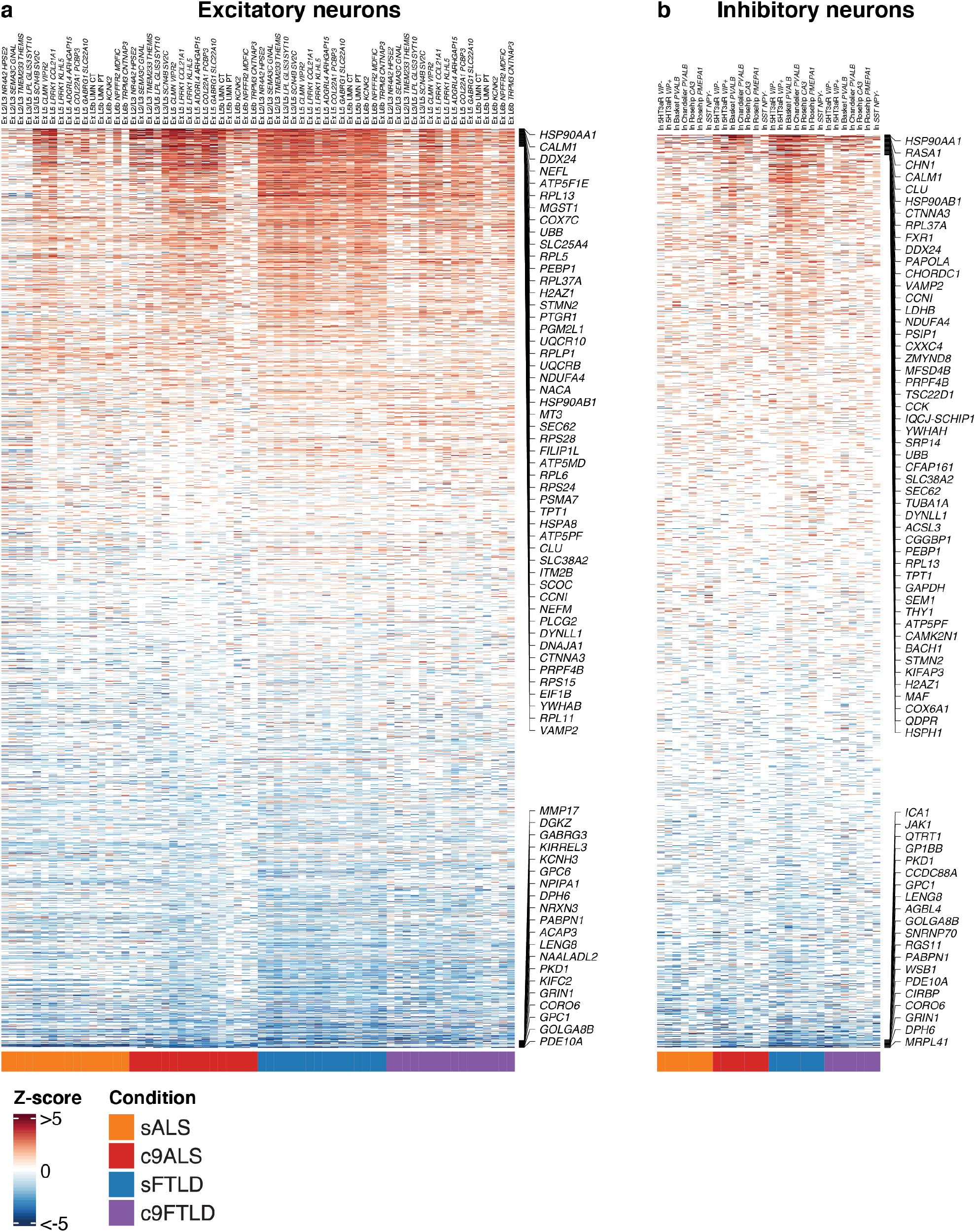
Top upregulated and downregulated DEGs in neuronal populations. Heatmaps of top 100 most upregulated and downregulated genes across cell types and disease groups for excitatory (**a**) and inhibitory (**b**) neuron populations (absolute log2-fold change Z-score > 1 and FDR-adjusted *p* < 0.005). Top 50 most upregulated and top 20 most downregulated across subtypes by mean row expression shown right of each heatmap. Many of the most upregulated DEGs observed in excitatory neurons were shared across subtypes and disease groups. Inhibitory neurons showed a more cell and phenotype specific response, but a subset of the common excitatory DEGs were also present and shared across inhibitory subtypes.

**Extended Data Figure 7.**
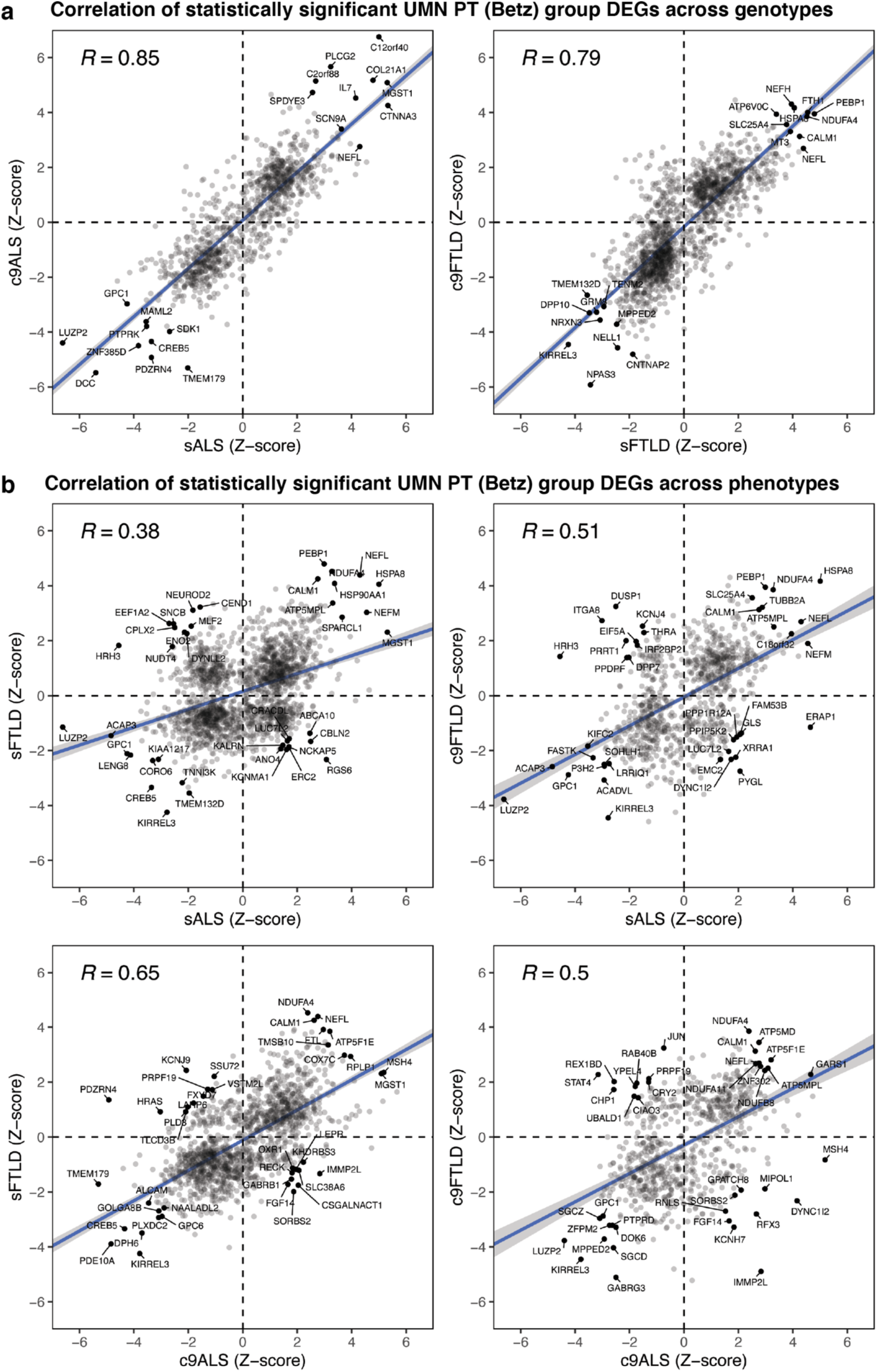
Comparison of Betz cell DEGs across genotypes and phenotypes. **a.** Scatter plots of cross-genotype comparison of statistically significant Betz cell DEGs within ALS (left) and FTLD (right) cohorts. Top 10 most up and down regulated genes labeled. **b.** Scatter plots of cross-phenotype comparison of statistically significant Betz cell DEGs between ALS and FTLD. Top 10 most up, down, and anti-directionally regulated genes labeled. FDR-adjusted *p* < 0.005 for all comparisons.

**Extended Data Figure 8.**
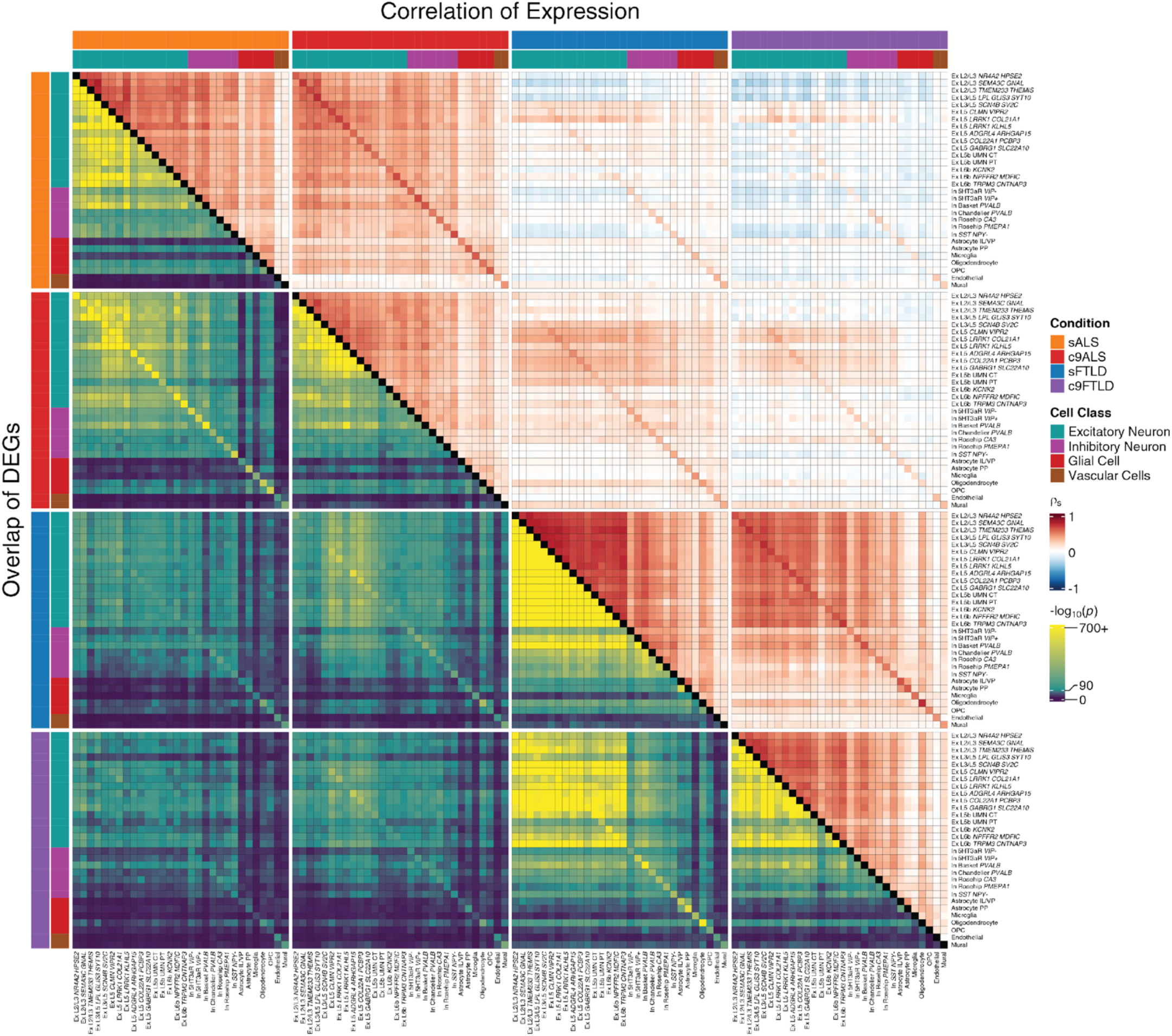
Heatmap of DEG correlation and overlap across cell types and disease groups. Upper diagonal: Spearman correlation of DEG *t*-statistic across all cell types and disease groups considering direction of dysregulation. Lower diagonal: Hypergeometric *p*-value of DEG overlap without considering direction of dysregulation. Absolute log2-fold change Z-score > 1 and FDR-adjusted *p* < 0.005 for all pairwise comparisons.

**Extended Data Figure 9.**
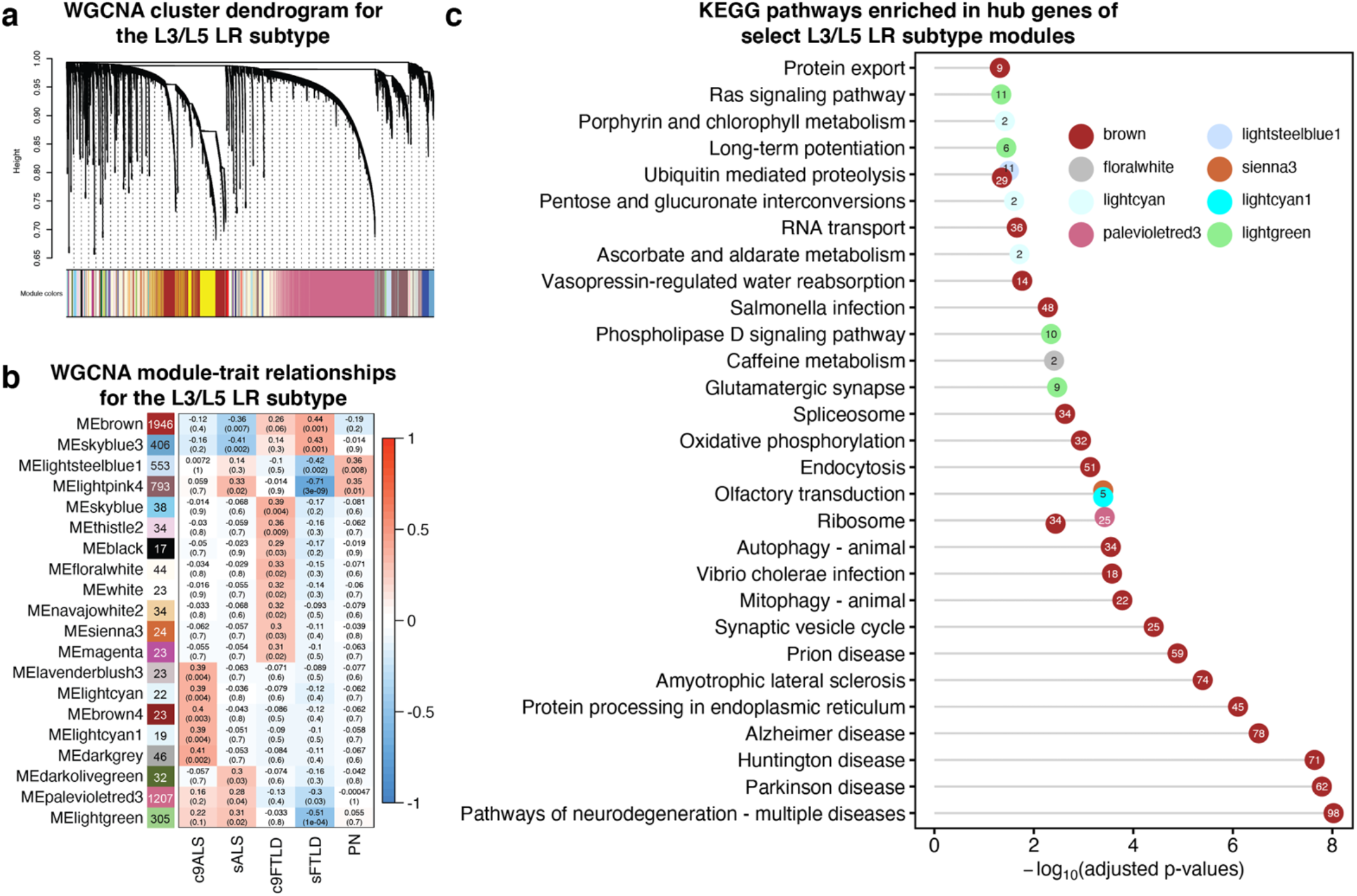
WGCNA analysis of Ex L3/L5 *SCN4B*/SC2C (L3/L5 LR) cluster in ALS and FTLD. **a.** Dendrogram showing result of unsupervised hierarchical clustering of modules identified by WGCNA. **b.** Pearson correlation of module eigengenes with disease groups. Modules significantly correlated with ALS and/or FTLD are shown, with the number of hub genes displayed on the right of each module’s name. **c.** Significant KEGG pathways enriched in hub genes of “brown”, “lightsteelblue1”, “floralwhite”, “sienna3”, “lightcyan”, “lightcyan1”, “palevioletred3”, and “lightgreen” modules. Numbers in each circle represent the number of hub genes in each pathway from the corresponding module.

**Extended Data Table 1.**
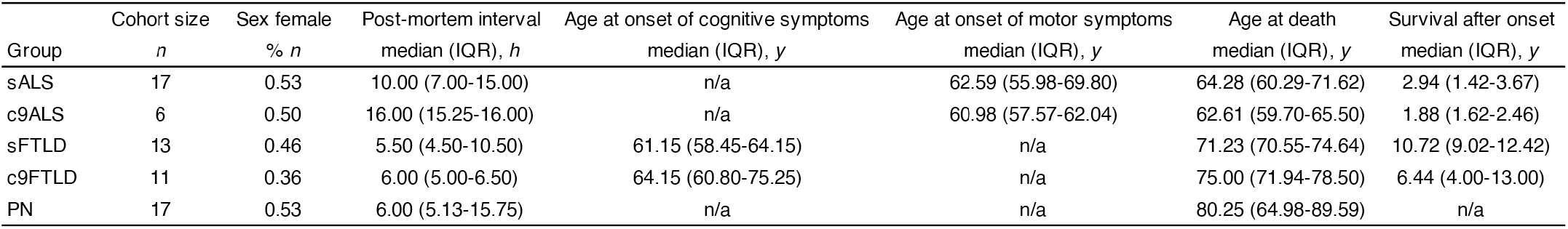
Summarized characteristics of profiled brain samples. Calculations include all individuals with available information: sex (64/64), post-mortem interval (PMI; 52/64), age at onset cognitive (22/24), age at onset motor (23/23), survival after onset (45/47). *n*: sample size; IQR: interquartile range; *h*: hours; *y*: years.

| Group | Cohort size<br>*n* | Sex female<br>% *n* | Post-mortem interval<br>median (IQR), *h* | Age at onset of cognitive symptoms<br>median (IQR), *y* | Age at onset of motor symptoms<br>median (IQR), *y* | Age at death<br>median (IQR), *y* | Survival after onset<br>median (IQR), *y* |
| --- | --- | --- | --- | --- | --- | --- | --- |
| sALS | 17 | 0.53 | 10.00 (7.00-15.00) | n/a | 62.59 (55.98-69.80) | 64.28 (60.29-71.62) | 2.94 (1.42-3.67) |
| c9ALS | 6 | 0.50 | 16.00 (15.25-16.00) | n/a | 60.98 (57.57-62.04) | 62.61 (59.70-65.50) | 1.88 (1.62-2.46) |
| sFTLD | 13 | 0.46 | 5.50 (4.50-10.50) | 61.15 (58.45-64.15) | n/a | 71.23 (70.55-74.64) | 10.72 (9.02-12.42) |
| c9FTLD | 11 | 0.36 | 6.00 (5.00-6.50) | 64.15 (60.80-75.25) | n/a | 75.00 (71.94-78.50) | 6.44 (4.00-13.00) |
| PN | 17 | 0.53 | 6.00 (5.13-15.75) | n/a | n/a | 80.25 (64.98-89.59) | n/a |

**Supplementary Data Table 1. Novel marker genes for annotated subtypes.** Table of top 100 subtype-specific marker genes identified by ACTIONet.

**Supplementary Data Table 2. Differential gene expression results - sALS.** Raw results of limma pseudo-bulk differential expression analysis for sporadic ALS cohort.

**Supplementary Data Table 3. Differential gene expression results – c9ALS.** Raw results of limma pseudo-bulk differential expression analysis for *C9orf72*-associated ALS cohort.

**Supplementary Data Table 4. Differential gene expression results - sFTLD.** Raw results of limma pseudo-bulk differential expression analysis for sporadic FTLD cohort.

**Supplementary Data Table 5. Differential gene expression results – c9FTLD.** Raw results of limma pseudo-bulk differential expression analysis for *C9orf72*-associated FTLD cohort.

**Supplementary Data Table 6. UMN PT hub genes identified by WGCNA.** Tables of hub genes corresponding to significant co-expression modules of UMN PT group.

**Supplementary Data Table 7. KEGG pathway enrichment for UMN PT hub genes.** Tables of pathway enrichment results for UMN PT group hub genes identified by WGCNA. Genes were compared against the Kyoto Encyclopedia of Genes and Genomes (KEGG) pathway database.

## Methods

### Human sample selection and preparation

Human tissue analysis was conducted as exempt human research, considering frozen post-mortem brain samples obtained from the Neuropathology Laboratory at the Mayo Clinic (Jacksonville, FL USA) were not specifically collected for this study. All cases were carefully analyzed by experienced and certified neuropathologists. TDP-43 pathology was confirmed in all ALS and FTLD samples based upon current consensus criteria, which investigates cortical and subcortical distribution of TDP-43 neuropathologic inclusions^114,115^. A section of the cerebellum was screened for *C9ORF72*-related pathology with P62 immunohistochemistry^116^. For all *C9orf72*-associated cases, repeat expansions were confirmed via Southern blot. Extent of upper and lower motor neuron involvement was investigated in all samples using Luxol fast blue and IBA-1 immunohistochemistry in the motor cortex, midbrain, and medulla. Motor neuron degeneration was confirmed in all ALS cases and found absent from all FTLD and control samples selected.

We selected a total of 64 age- and sex-matched individuals with sporadic ALS (sALS; *n*=17), *C9orf72*-associated ALS (c9ALS; *n*=6), sporadic FTLD (sFTLD; *n*=13), *C9orf72*-associated FTLD (c9FTLD; *n*=11), or found pathologically normal (*n*=17). Each cohort included similar numbers of male and female samples (sALS 8:9; c9ALS 3:3; sFTLD 7:6; c9FTLD 7:4; PN 8:9). Each *C9orf72*-associated disease cohort included patients with a positive family history of either ALS or FTLD. All other disease cases selected were considered sporadic, which is representative of the majority of patients: no family history and no defined genetic risk factor (no mutation in *SOD1*, *TARDBP*, *FUS*, *C9orf72*, *NEK1*, *GRN*, *MAPT*, or *TBK1*). For each brain selected, approximately 300mg of the primary motor cortex was dissected by the Mayo Clinic Neuropathology Laboratory.

### Isolation of nuclei from post-mortem frozen human brain tissue for single nuclear RNA-sequencing

Nuclei isolation protocol was adapted from Lee et al.^117^ All procedures were performed on ice. Tissue was homogenized in 700 μL of homogenization buffer (320 mM sucrose, 5 mM CaCl_2_, 3 mM Mg(CH_3_COO)_2_, 10 mM Tris HCl [pH 7.8], 0.1 mM EDTA [pH 8.0], 0.1% NP-40, 1 mM β-mercaptoethanol, and 0.4 U/μL SUPERaseIn RNase Inhibitor (ThermoFisher Scientific, Waltham MA) with a 2 mL KIMBLE Dounce tissue grinder (MilliporeSigma, Burlington MA) using 10 strokes with loose pestle followed by 10 strokes with tight pestle. From the mouse tissue samples the entire striatum was homogenized. From the human tissue samples 100 mg of grey matter was sectioned and homogenized. Homogenized tissue was filtered through a 40 μm cell strainer and mixed with 450 μL of working solution (50% OptiPrep density gradient medium (MilliporeSigma, Burlington MA), 5 mM CaCl_2_, 3 mM Mg(CH_3_COO)_2_, 10 mM Tris HCl [pH 7.8], 0.1 mM EDTA [pH 8.0], and 1 mM β-mercaptoethanol). The mixture was then slowly pipetted onto the top of an OptiPrep density gradient containing 750 μL of 30% OptiPrep Solution (134 mM sucrose, 5 mM CaCl_2_, 3 mM Mg(CH_3_COO)_2_, 10 mM Tris HCl [pH 7.8], 0.1 mM EDTA [pH 8.0], 1 mM β-mercaptoethanol, 0.04% NP-40, and 0.17 U/μL SUPERase In RNase Inhibitor) on top of 300 μL of 40% OptiPrep Solution (96 mM sucrose, 5 mM CaCl_2_, 3 mM Mg(CH_3_COO)_2_, 10 mM Tris HCl [pH 7.8], 0.1 mM EDTA [pH 8.0], 1 mM β-mercaptoethanol, 0.03% NP-40, and 0.12 U/μL SUPERase In RNase Inhibitor) inside a Sorenson Dolphin microcentrifuge tube (MilliporeSigma, Burlington MA). Nuclei were pelleted at the interface of the OptiPrep density gradient by centrifugation at 10,000 × *g* for 5 min at 4°C using a fixed angle rotor (FA-45-24-11-Kit). The nuclear pellet was collected by aspirating ~100μL from the interface and transferring to a 2.5 mL Eppendorf tube. The pellet was washed with 2% BSA (in 1x PBS) containing 0.12 U/μL SUPERase In RNase Inhibitor. The nuclei were pelleted by centrifugation at 300 × *g* for 3 min at 4°C using a swing-bucket rotor (S-24-11-AT). Nuclei were washed three times with 2% BSA and centrifuged under the same conditions. The nuclear pellet was resuspended in 100 μL of 2% BSA.

### Sequencing data preprocessing and analysis

Droplet-based snRNA sequencing libraries were prepared using the Chromium Single Cell 3’ Reagent Kit v3 (10x Genomics, Pleasanton CA) according to the manufacturer’s protocol and sequenced on a NovaSeq 6000 at the Broad Institute Genomics Platform. FASTQ files were aligned to the pre-mRNA annotated human reference genome GRCh38. Cell Ranger v4.0 (10x Genomics, Pleasanton CA) was used for genome alignment and feature-barcode matrix generation.

We used the ACTIONet^29,118^ and scran^119^ R packages to normalize, batch correct, and cluster single-cell gene counts. A curated set of known cell type-specific markers was used to annotate individual cells with their expected cell type and assign a confidence score to each annotation. We removed cells with high mitochondrial RNA content, abnormally low or high RNA content (relative to the distribution of its specific cluster), ambiguous overlapping profiles resembling dissimilar cell types (generally corresponding to doublet nuclei), and cells corresponding to graph nodes with a low *k*-core or low centrality in the network (generally corresponding to high ambient RNA content or doublet nuclei).

Cell type-specific pseudo-bulk differential gene expression (DGE) analysis was performed using ACTIONet and limma^120^ for sufficiently abundant cell types using age, sex, and disease group as design covariates and gene-wise single-cell-level variance as weights for the linear model. We found Braak stage to be a poor predictor of gene expression and omitted it from the design formula. Genes were considered differentially expressed if they had an FDR-corrected p-value < 0.005 and an absolute log2-fold change > 1 standard deviation for that cell type relative to the pathologically normal control group. To ensure that DGE results were reproducible and robust to differences in cell type abundance, we sampled with replacement equal numbers of individuals and cells per individual for each cell type and repeated the pseudo-bulk analysis. Lastly, we repeated the analyses using DESeq2^121^ as the model-fitting algorithm in lieu of limma to ensure replicability across methods. In all cases DGE results were consistent and we used the pseudo-bulk limma results for all downstream analyses. Pathway analysis was performed using the g:Profiler^122^ R package.

### WGCNA

We used R package WGCNA^70–72^ to perform the weighted correlation network analysis on pseudo-bulk expression profiles. A signed network was constructed with the UMN PT and L3/L5 LR clusters shown in the ACTIONet plot of excitatory neurons (**Fig. 1b**). The co-expression similarity was raised to the power ß = 8 to arrive at the network adjacency (R^2^ is 0.8 when ß = 8). The modules were identified using the R function *blockwiseConsensusModules* with minModuleSize = 30. Hub genes (genes with highest module membership) in each consensus module were identified using the R function *signedKME^73^*. Pathway analysis was performed using the R package gprofiler2^122^ considering only protein-coding hub genes with kME value > 0.6.

### Immunofluorescent labeling and confocal microscopy

Patient tissues were embedded in Allan Scientific™ Neg-50™ Frozen Section Medium (Thermo Scientific #6502), cryosectioned to 20μm thick, and transferred to glass slides (VWR #48311-703). Next, the tissues were subject to 100% ice-cold acetone (Sigma-Aldrich #320110) for 10 minutes followed by 15 minutes of tris-buffered saline with 0.25% Triton X-100 then blocked for 1 hour in tris-buffered saline with Tween 20 solution at 0.05% (BioRad #161-0781) supplemented with sterile 2% Normal Donkey serum (Abcam #ab138579) and 0.1% fish gelatin (Sigma-Aldrich #G7765). Next, the slides were incubated overnight at 4 degrees Celsius in a slide humidity chamber with primary antibodies POU3F1 (Millipore Sigma MABN738) and TDP-43 (ProteinTech 10782-2-AP) or CRYM (Abcam ab220085). Next, the slides were washed and incubated with Donkey anti-Mouse IgG (H+L) 488nm and Donkey anti-Rabbit IgG (H+L) 546nm (ThermoFisher #A-21202, A-10040, respectively) for Figure 5a. The slides were washed and incubated with Donkey anti-Mouse IgG (H+L) 488nm and Donkey anti-Rabbit IgG (H+L) 647nm (ThermoFisher #A-21202, #A-31573, respectively) for Figure 5b. Following additional washes, a 10 mg/mL stock of Hoechst 33342, Trihydrochloride, Trihydrate (ThermoFisher #H1399) was used at 1μL per 10mL of washing solution for 10 minutes. Next, the tissues were treated with a solution containing TrueBlack (Biotium) at 50μL per 1 mL of 70% ethanol for 10 seconds, then washed with tris-buffered saline solution (no detergent). Tissues were then mounted using ProLong™ Gold Antifade Mountant (ThermoFisher #P36930). Mounted slides were imaged on a Zeiss Observer.Z1 LSM 700 confocal microscope (Carl Zeiss AG, Oberkochen, Germany) using a EC Plan-Neofluar 40X/1.30 Oil DIC M27 objective. Z-stack images were max-projected using Fiji^123^.

## Acknowledgements

The work was supported by grants from the National Institutes of Health (NIH), Mitsubishi Tanabe Pharma Holdings America, Inc., the JPB Foundation, the Robert Packard Center for ALS Research at Johns Hopkins, the LiveLikeLou Foundation, the Gerstner Family Foundation, the Mayo Clinic Center for Individualized Medicine, and the Cure Alzheimer’s Fund. NIH NS100802 to M.H.; NIH AG067151 to V.V.B.; NIH AG054012, AG058002, AG062377, NS110453, NS115064, AG067151, AG062335, MH109978, MH119509, HG008155 to M.K.; NIH 5T32EB019940-05 to S.S.P.; NIH NS110994 to M.V.B.; NIH AG37491 to K.A.J.. H.L. was supported by a postdoctoral fellowship from the JPB Foundation, and B.E.F. was supported by a postdoctoral fellowship from the Hereditary Disease Foundation. Brain samples were provided by the Neuropathology Laboratory at the Mayo Clinic (Jacksonville, FL USA). We would like to thank all study participants for donating their brain to research, and Sami Barmada, M.D., Ph.D., for his professional input.

## Data Availability

Raw and processed sequencing data, including annotated count matrices, associated with Figure 1 will be available on NCBI GEO upon publication.

## Code Availability

Code used in this study is available upon reasonable request.

## Author Contributions

S.S.P., H.L., and J.M. carried out single-cell profiling. S.S.P., H.L., and S.M. carried out computational analyses. B.E.F. carried out experimental validations. L.J.P., M.E.G., E.E.-C., M.D.-H., M.V.B., C.P., R.R., B.O., J.S.S., R.C.P., N.R.G.-R., B.F.B., D.S.K., K.A.J., M.D., M.E.M., D.W.D., and V.V.B. provided biological samples and phenotype/genotype information. M.H., M.K., and V.V.B. co-directed the work. S.S.P., H.L., B.E.F., M.H., V.V.B., and M.K. wrote the manuscript with input from all authors.

## Supplementary Information

Supplementary information will be made available at http://compbio.mit.edu/scALS/.

